# The AP-1 complex is essential for fungal growth via its role in secretory vesicle polar sorting, endosomal recycling and cytoskeleton organization

**DOI:** 10.1101/294223

**Authors:** Olga Martzoukou, George Diallinas, Sotiris Amillis

## Abstract

The AP-1 complex is essential for membrane protein traffic via its role in the pinching-off and sorting of secretory vesicles from the trans-Golgi and/or endosomes. While its essentiality is undisputed in metazoa, its role in model simpler eukaryotes seems less clear. Here we dissect the role of AP-1 in the filamentous fungus *Aspergillus nidulans* and show that it is absolutely essential for growth due to its role in clathrin-dependent maintenance of polar traffic of specific membrane cargoes towards the apex of growing hyphae. We provide evidence that AP-1 is involved in both anterograde sorting of RabE^Rab11^-labeled secretory vesicles and RabA/B^Rab5^- dependent endosome recycling. Additionally, AP-1 is shown to be critical for microtubule and septin organization, further rationalizing its essentiality in cells that face the challenge of cytoskeleton-dependent polarized cargo traffic. This work also opens a novel issue on how non-polar cargoes, such as transporters, are sorted to the eukaryotic plasma membrane.

## Introduction

All eukaryotic cells face the challenge of topological sorting of their biomolecules in their proper subcellular destinations. In particular, newly synthesized membrane proteins, which are translationally translocated into the membrane of the ER, follow complex, dynamic, and often overlapping, trafficking routes, embedded in the lipid bilayer of „secretory‟ vesicles, to be sorted to their final target membrane (Feyder et al., 2015; Viotti 2016). In vesicular membrane trafficking, the nature of the protein cargo and relevant adaptor proteins play central roles in deciding the routes followed and the final destination of cargoes. Despite the emerging evidence of alternative or non-conventional trafficking routes, cargo passage through a continuously maturing early-to-late Golgi is considered to be part of the major mechanism and the most critical step in membrane protein sorting. Following exit from the trans-Golgi network (TGN, also known as late-Golgi), cargoes packed in distinct vesicles travel to their final destination, which in most cases is the plasma membrane or the vacuole. This anterograde vesicular movement can be direct or via the endosomal compartment, and in any case assisted by motor proteins and the cytoskeleton (Bard and Malhotra 2006; Cai et al., 2007; Hunt and Stephens 2011; Anitei and Hoflack 2011; Guo et al., 2014). Membrane protein cargoes at the level of late Golgi can also follow the opposite route, getting sorted into retrograde vesicles, recycling back to an earlier compartment. Acquiring a “ticket” for a specific route implicates adaptors and accessory proteins, several of which are also associated with clathrin (Nakatsu and Ohno 2003; Robinson 2004; 2015).

Apart from the COPI and COPII vesicle coat adaptors that mediate traffic between the ER and the early Golgi compartment (Lee et al., 2004; Zanetti et al., 2011), of particular importance are the heterotetrameric AP (formally named after Assembly Polypeptide and later as Adaptor Protein) complexes, comprising of two large subunits (also called adaptins; β-adaptin and γ- or α-adaptin), together with a medium-sized (μ) and a small (σ) subunit (Robinson 2004; 2015). Other adaptors, some of which display similarity to AP subunits, like the GGAs, epsin-related proteins, or components of the exomer and retromer complexes are also critical for the sorting of specific cargoes (Bonifacino 2004; 2014; Robinson 2015; Spang 2015; Anton et al., 2018). Importantly, the various cargo sorting routes are often overlapping and might share common adaptors (Hoya et al., 2017). Among the major AP complexes (Bonifacino 2014; Nakatsu et al., 2014), AP-1 and AP-2, which in most cells work to propel vesicle formation through recruitment of clathrin, are the most critical for cell homeostasis and function (Robinson 2004; 2015). In brief, AP-2 is involved in vesicle budding for protein endocytosis from the PM, whereas AP-1 is involved in vesicle pinching-off from the TGN and/or endosomal compartments, although in the latter case it is still under debate whether secretory vesicles derive from the TGN, from the endosome, or from both (Nakatsu et al., 2014; Robinson 2015). AP-1 was also shown to be responsible for retrograde transport from early endosomes in both yeast and mammalian cells, but also guiding recycling pathways from the endosome to the plasma membrane in yeast (Spang 2015). The undisputed essentiality of AP-1 and AP-2 in mammalians cells is however less obvious in simple unicellular eukaryotes, such as the yeasts *Saccharomyces cerevisiae* or *Schizosaccharomyces pombe*, where null mutants in the genes encoding AP subunits are viable, with only relatively minor growth or morphological defects (Meyer et al., 2000; Valdivia *et al.* 2002; Ma et al., 2009; Yu et al., 2013; Arcones et al., 2016). In sharp contrast, the growth of AP-1 and AP-2 null mutants in the filamentous fungus *Aspergillus nidulans* is severally arrested after spore germination (Martzoukou et al., 2017 and results presented herein), reflecting blocks in essential cellular processes, probably similar to mammalian cells.

In the recent years, *A. nidulans* is proving to be a powerful emerging system for studying membrane cargo traffic (Momany 2002; Taheri-Talesh et al., 2008; Steinberg et al., 2017). This is not only due to its powerful classical and reverse genetic tools, but also due to its specific cellular characteristics and way of growth. *A. nidulans* is made of long cellular compartments (hyphae), characterized by polarized growth, in a process starting with an initial establishment of a growth site, followed by polarity maintenance and cell extension through the regulated continuous supply of vesicles towards the apex. A vesicle sorting terminal at the hyphal apex, termed Spitzenkörper (Spk), is thought to generate an exocytosis gradient, which when coupled with endocytosis from a specific hotspot behind the site of growth, termed endocytic collar, is able to sustain apical growth (Penalva 2015; Schultzhaus and Shaw 2015; Pantazopoulou 2016; Steinberg et al., 2017). Apical trafficking of cargoes, traveling from the endoplasmic reticulum (ER) through the different stages of early (cis-) and late (trans-) Golgi towards their final destination, and apical cargo endocytosis/recycling, are essential for growth, as null mutations blocking either Golgi function, microtubule organization or apical cargo recycling are lethal or severely deleterious (Fischer et al., 2008; Takeshita and Fischer 2011; Penalva 2015; Pantazopoulou 2016; Steinberg et al., 2017). Curiously, the role of AP complexes in *A. nidulans* or any other filamentous fungus, has not been studied, with the exception of our recent work on AP-2 (Martzoukou et al., 2017). In that study we showed that the AP-2 of *A. nidulans* has a rather surprising clathrin-independent essential role in polarity maintenance and growth, related to the endocytosis of specific polarized cargoes involved in apical lipid and cell wall composition maintenance. This was in line with the observation that AP-2 β subunit (β2) lacks the ability to bind clathrin, which itself has been shown to be essential for the endocytosis of distinct cargoes, as for example various transporters (Martzoukou et al., 2017; Schultzhaus et al., 2017). In the current study, we focus on the role of the AP-1 complex in cargo trafficking in *A. nidulans*. We provide evidence that AP-1 is essential for fungal polar growth via its dynamic role in post-Golgi secretory vesicle polar sorting, proper microtubule organization and endosome recycling.

## Results

### The AP-1 complex localizes polarly in distinct cytoplasmic structures and is essential for growth

In *A. nidulans*, the AP-1 adaptor complex is encoded by the genes AN7682 (*ap-1*^σ^), AN4207 (*ap-1*^γ^), AN3029 (*ap-1*^β^) and AN8795 (*ap-1*^μ^). In a previous study a knockout of the gene encoding the AP-1 σ subunit proved lethal, therefore we employed a conditional knock-down strain (Martzoukou et al., 2017), using the thiamine-repressible promoter *thiA*_*p*_ (Apostolaki et al., 2012). The phenotypic analysis of this strain showed that repression of *ap1*^σ^ results in severely retarded colony growth, reflected at the microscopic level in wider and shorter hyphae with increased numbers of side branches and septa. Figure 1A and 1B highlight these results, further showing that *thiA*_*p*_–dependent full repression of not only *ap-1*^σ^, but also *ap-1*^β^ and *ap-1*^μ^, results in lack of growth. Notably, besides increased numbers of side branches and septa, staining level and cortical localization of calcofluor white are modified upon repression of AP-1^σ^, suggesting altered chitin deposition (Figure 1B). Given that the genetic disruption of three AP-1 subunits appears to affect growth in *A. nidulans* similarly and also that the inactivation of any subunit has been reported to disrupt the full complex in other organisms (Robinson 2015 and refs therein), the AP-1^σ^ subunit was chosen to further investigate the role of the AP-1 complex in intracellular cargo trafficking pathways.

**Figure 1:**
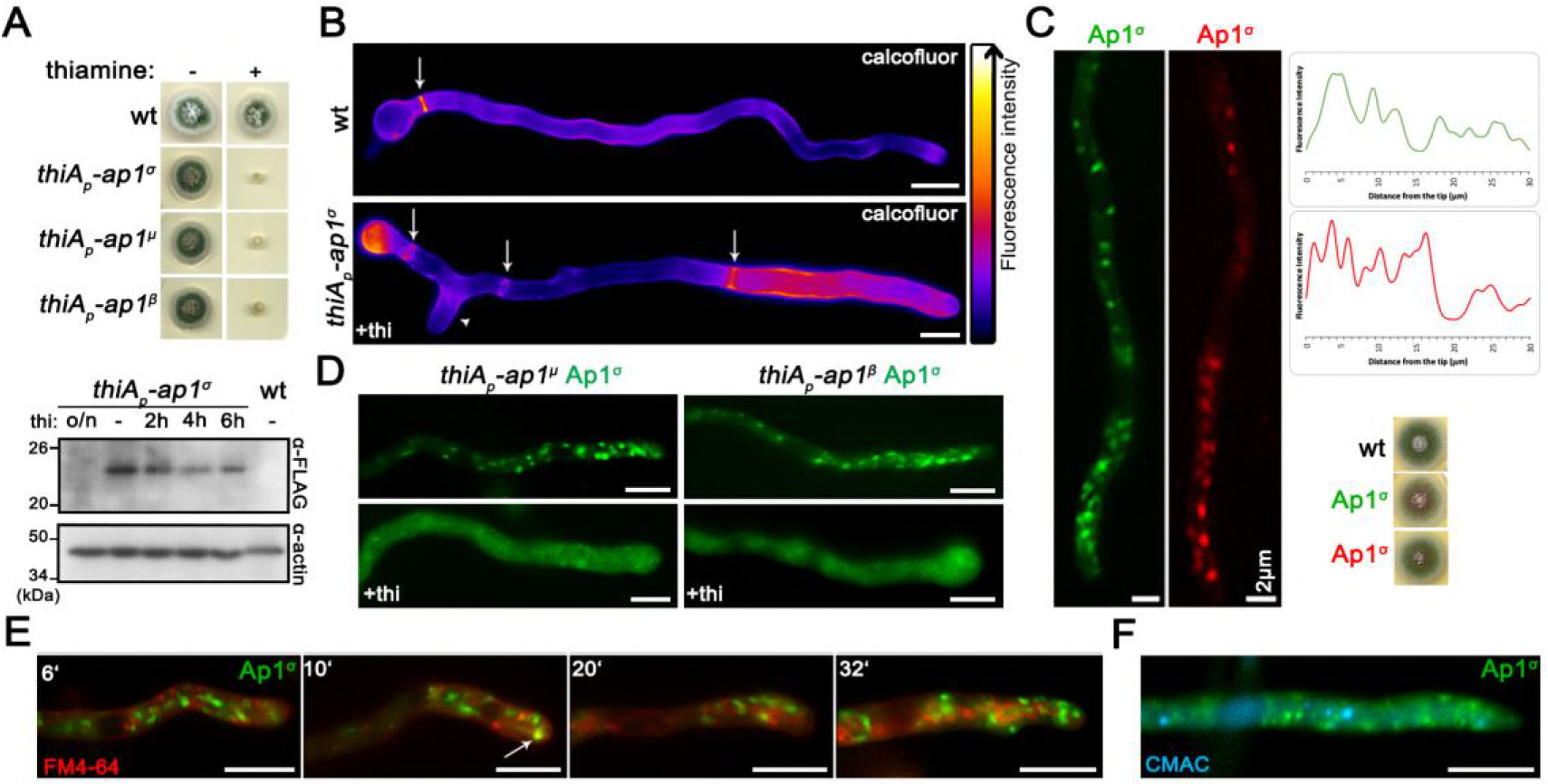
The AP-1 complex localizes in distinct polarly distributed structures and is essential for growth. (**A**) Upper panel: Growth of isogenic strains carrying thiamine-repressible alleles of *ap1^σ^*, *ap1^μ^* and *ap1^β^* (*thiA_p_-ap1^σ^, thiA_p_-ap1^μ^* and *thiA_p_-ap1^β^*) compared to wild-type (*wt*) in the absence (-) or presence (+) of thiamine. Lower panel: Western blot analysis comparing protein levels of FLAG-Ap1^σ^ in the absence (0h) or presence of thiamine, added for 2, 4, 6 or 16h (overnight culture, o/n). *wt* is a standard wild-type strain (untagged *ap1^σ^*) which is included as a control for the specificity of the α-FLAG antibody. Equal loading is indicated by actin levels. (**B**) Microscopic morphology of hyphae in a strain repressed for *ap1^σ^* expression (+thi, lower panel) compared to *wt* (upper panel) stained with calcofluor white. Septal rings and side branches are indicated by arrows and arrowheads. Notice the differences in the calcofluor deposition at the hyphal head, tip and the sub apical segment (Lookup table [LUT] fire [ImageJ, National Institutes of Health]) (**C**) Subcellular localization of Ap1^σ^-GFP and Ap1^σ^-mRFP in isogenic strains and relative quantitative analysis of fluorescence intensity (right upper panel), highlighting the polar distribution of Ap1^σ^. Growth tests showing that the tagged versions of Ap1^σ^ are functional (right lower panel). (**D**) Subcellular localization of Ap1^σ^-GFP in isogenic strains carrying thiamine-repressible alleles of *ap1^μ^* (left panels) or *ap1^β^* (right panels) in the absence (upper panels) or presence of thiamine (+thi, o/n). Notice that repression of expression of either the μ or the β subunit leads to diffuse cytoplasmic fluorescent of Ap1^σ^. (**E**) Subcellular localization of Ap1^σ^-GFP in the presence of FM4-64, which labels dynamically endocytic steps (PM, early endosomes, late endosomes/vacuoles). Notice that Ap1^σ^- GFP structures do not co-localize with FM4-64, except a few cases observed in the sub-apical region (indicated with an arrow at the 10 min picture). (**F**) Subcellular localization of Ap1^σ^-GFP in the presence of the vacuolar stain 7-amino-4- chloromethylcoumarin (Blue CMAC). No Ap1^σ^-GFP/CMAC co-localization is observed. Unless otherwise stated, scale bars represent 5 μm.

Figure 1C shows that expression of functional GFP- or mRFP-tagged AP-1^σ^ has distinct localization in cytoplasmic puncta, the motility of which resembles a Brownian motion, being more abundant in the apical region of hyphae and apparently absent from the Spk. The distinct localization of AP-1^σ^ localization, which resembles the distribution of Golgi markers (see later), is lost and replaced by a fluorescence cytoplasmic haze when the β or μ subunits are knocked-down (Figure 1D). Noticeably, the majority of these foci are not stained by FM4-64 or CMAC (Figure 1E, 1F), strongly suggesting that they are distinct from endosomes and vacuoles.

### Knockdown of AP-1 affects the localization of polarly localized cargoes

As mentioned in the Introduction, polarized growth of fungal hyphae is sustained by the continuous delivery of cargo-containing secretory vesicles (SV) towards the hyphal apex and accumulation at the Spk before fusion with the plasma membrane (PM). Once localized in the PM at the hyphal apex, several cargoes diffuse laterally and get recycled through the actin-patch enriched subapical regions of the endocytic collar, balancing exocytosis with endocytosis (Harris 2005; Steinberg 2007; Berepiki et al., 2011; Takeshita et al., 2014; Penalva et al., 2017; Steinberg et al., 2017; Zhou et al., 2018). In order to study the potential implications of the AP-1 complex in these processes, we monitored the localization of specific established apical and collar markers in conditions where the *ap-1*^σ^ expression has been fully repressed. These markers include the secretory v-SNARE SynA and t-SNARE SsoA (Taheri-Talesh et al., 2008), the phospholipid flippases DnfA and DnfB that partially localize in the Spk (Schultzhaus et al., 2015), the class III chitin synthase ChsB known to play a key role in hyphal tip growth and cell wall integrity maintenance (Yanai et al., 1994; Takeshita et al., 2015), the tropomyosin TpmA decorating actin at the Spk and on actin cables at the hyphal tip (Taheri-Talesh et al., 2008), and finally the endocytic patch specific marker AbpA marking the sites of actin polymerization (Araujo-Bazán et al., 2008), along with the endocytic markers SlaB and SagA (Araujo-Bazán et al., 2008; Hervás-Aguilar and Peñalva, 2010; Karachaliou et al., 2013). Additionally, we also tested the localization of the UapA xanthine-uric acid transporter, for which our previous work suggested that it is not affected by the loss of function of the AP-1 complex (Martzoukou et al., 2017).

Figure 2 highlights our results, which show that the localization of all markers tested is affected in the absence of AP-1^σ^, with the only clear exception being the plasma membrane transporter UapA. Additionally, the general positioning of nuclei also appears unaffected as indicated by labeled Histone 1 (Nayak et al., 2010). Of the markers tested, SynA, DnfA, DnfB and ChsB lose significantly their polar distribution and do not seem to properly reach the Spk, concomitant with their increased presence in distinct, rather static, cytoplasmic puncta of various sizes. The non-polar distribution of SsoA is generally conserved, but its cortical positioning is reduced and replaced by numerous cytoplasmic puncta. All collar-associated markers (SagA, SlaB and AbpA) appear “moved” in an acropetal manner towards the hyphal tip. TpmA has also lost its proper localization at the hyphal tip, suggesting defective stabilization of actin filaments at the level of the Spk (Bergs et al., 2016). For relative quantification of fluorescence intensity see also Figure 2 Supplement 1.

**Figure 2:**
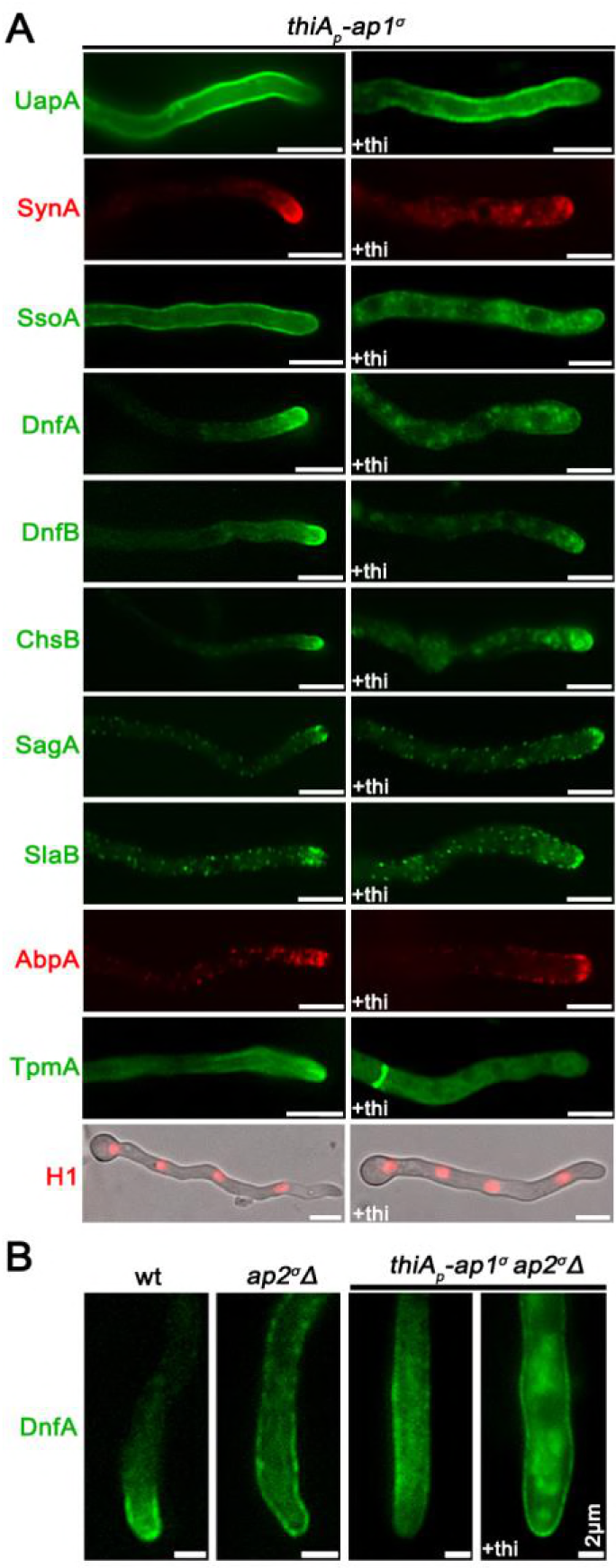
Lack of expression of AP-1 affects the topology of polar cargoes. (**A**) Comparison of the cellular localization of specific GFP- or mRFP/mCherry-tagged protein cargoes under conditions where *ap1^σ^* is expressed (left panel) or fully repressed by thiamine (right panel, +thi). The cargoes tested are the UapA transporter, the SNAREs SynA and SsoA, phospholipid flippases DnfA and DnfB, chitin synthase ChsB, endocytic markers SagA and SlaB, the actin-polymerization marker AbpA, tropomyosin TpmA, and histone H1 (i.e. nuclei). Notice that when *ap1^σ^* is fully repressed polar apical cargoes are de-polarized and mark numerous relatively static cytoplasmic puncta. (**B**) Localization of DnfA-GFP in strains carrying the *ap2^σ^*Δ null allele, or the *ap2^σ^*Δ null allele together with the repressible *thiA_p_-ap1^σ^* allele, or an isogenic wild-type control (*wt*: *ap2^σ^*^+^ *ap1^σ^*^+^). Notice that loss of polar distribution due to defective apical endocytosis observed the *ap2*^σ^Δ strains (Martzoukou et al., 2017) persists when AP-1^σ^ is also repressed, indicating that in the latter case the majority of accumulating internal structures are due to problematic exocytosis of DnfA. Unless otherwise stated, scale bars represent 5 μm.

Previous studies have shown that mutations preventing endocytosis of polar markers result in a uniform rather than polarized distribution of cargoes (Schultzhaus et al., 2015; Schultzhaus and Shaw, 2016; Martzoukou et al., 2017). In contrast, when exocytosis is impaired due to the absence of clathrin, several cargoes show predominantly non-cortical cytoplasmic localization (Martzoukou et al., 2017). Thus, our present observations strongly suggest that secretion and/or recycling is the process blocked in the absence of the AP-1 complex, while endocytosis remains functional. The latter is further supported by the fact that repression of AP-1 in the absence of a functional AP-2 complex, results in significant apparent cortical retention of specific cargoes, such as DnfA, despite the concurrent subapical accumulation of cytoplasmic DnfA-labeled structures (Figure 2B), which notably do not co-localize with endocytic membranes stained by FM4-64 (Figure 2 Supplement 2).

### AP-1 associates transiently with the trans-Golgi

In *A. nidulans*, the process of maturation of Golgi has been extensively studied (Peñalva 2010; Pantazopoulou 2016; Steinberg et al., 2017). The markers syntaxin SedV^Sed5^ and the human oxysterol-binding protein PH domain (PH^OSBP^) are well-established markers to follow the dynamics of early/cis-Golgi (Pinar et al., 2013) and late/trans-Golgi compartments (Pantazopoulou and Peñalva, 2009), respectively. Here we examined the possible association of the AP-1 complex with Golgi compartments using these markers. Figure 3A shows that AP-1^σ^ shows no co-localization, despite some topological proximity, with the early-Golgi, although in some cases it orbits around SedV marker (see also Video 1). Contrastingly, most AP-1^σ^ labeled structures show a significant degree of apparent association with PH^OSBP^, which suggests AP-1 partially co-localizes with the late-Golgi (Figure 3B, see also Video 2). Notably, the degree of association of AP-1 with PH^OSBP^ has a transient character, as seen by the apparent progressive loss of co-localization. The increased association of AP-1^σ^ with late-Golgi is further supported by the effect of Brefeldin A, which leads to transient Golgi collapse in aggregated bodies, several of which included the AP-1 marker (Figure 3C). Thermosensitive mutations in SedV (SedV-R258G) or the regulatory GTPase RabO^Rab1^ (RabO-A136D) are known to lead to early- or early/late-Golgi disorganization upon shift to the restrictive temperature (Pinar et al., 2013). These mutations led to AP-1^σ^ subcellular distribution modification, further supporting the association of AP-1 with late, but not with early Golgi. In particular, in SedV-R258G, AP-1^σ^ localization was less affected, whereas in RabO-A136D AP-1^σ^ fluorescence was significantly de-localized from distinct puncta to a cytoplasmic haze (Figure 3D). Finally, knockdown of RabC^Rab6^, another small GTPase responsible for Golgi network organization, also results in smaller AP-1^σ^ foci (Figure 3D), resembling the fragmented Golgi equivalents observed for PH^OSBP^ in a *rabC*Δ genetic background (Pantazopoulou and Peñalva, 2011).

**Figure 3:**
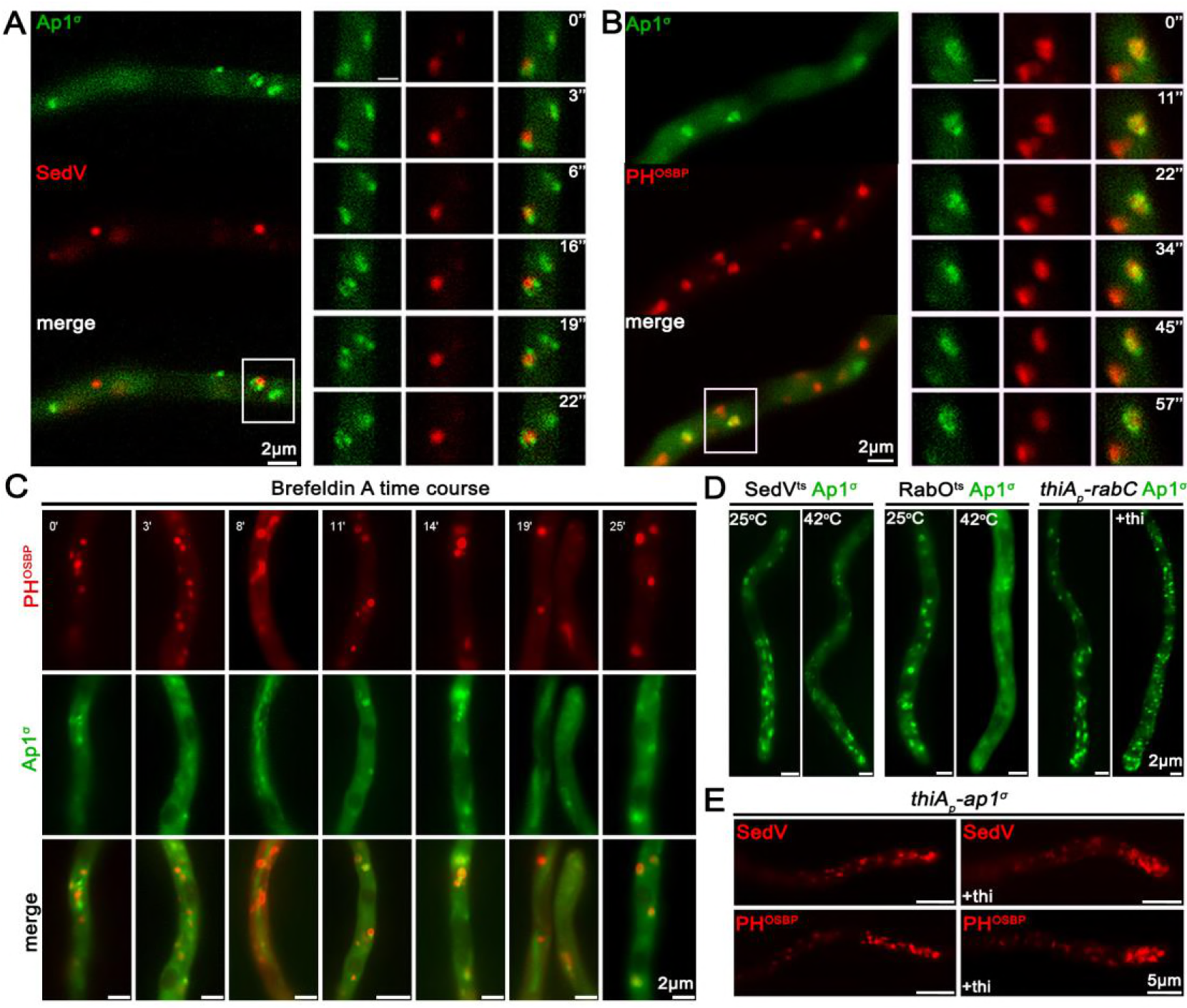
AP-1 associates transiently with the late-Golgi. (**A, B**) Subcellular localization of Ap1^σ^-GFP relative to *cis-* (SedV-mCherry) and *trans*-Golgi (PH^OSBP^-mRFP) markers. Notice that Ap1^σ^ co-localizes significantly with the *trans*-Golgi marker PH^OSBP^, but not with the *cis*-Golgi marker SedV. This co-localization is dynamic and transient, as shown in selected time lapse images on the right panels (see also relevant Videos 1 and 2). (**C**) Subcellular localization of Ap1^σ^ and PH^OSBP^ in the presence of the inhibitor Brefeldin A, showing that a fraction of Brefeldin bodies (i.e. collapsed Golgi membranes) includes both markers, further supporting a transient AP-1/late Golgi association. (**D**) Subcellular localization of Ap1^σ^ in SedV^ts^ or RabO^ts^ thermosensitive mutants or a strain carrying a repressible *rabC* allele. These strains are used as tools for transiently blocking proper Golgi function. Notice that at the restrictive temperature (42 °C) Ap1^σ^ fluorescence becomes increasingly diffuse mostly in the RabO^ts^ mutant, whereas under RabC repressed conditions small Ap1^σ^–labeled puncta increase in number. These results are compatible with the notion that AP-1 proper localization necessitates wild-type Golgi dynamics. (**E**) Distribution of early and late Golgi markers SedV and PH^OSBP^ relative to *ap1^σ^* expression or repression (+thi). Notice the effect of accumulation of Golgi towards the hyphal apex under repressed conditions. Unless otherwise stated, scale bars represent 5 μm.

Importantly, knockdown of AP-1 had a moderate but detectable effect on the overall picture of early- or late-Golgi markers, which in this case seem re-located in the subapical region of the hypha, thus showing increased polarization (Figure 3E; Figure 3 Supplement 1). The significance of this observation is discussed later.

### AP-1 associates with Clathrin via specific C-terminal motifs

AP-1 and AP-2 association with clathrin is considered as a key interaction mediating the recognition of cargo prior to clathrin cage assembly in metazoa (Robinson, 2015). Clathrin-binding motifs, or boxes, have been identified in the hinge regions of the β subunits (Dell‟Angelica et al., 1998). In *A. nidulans*, clathrin light and heavy chains have been recently visualized (Martzoukou et al., 2017; Schultzhaus et al., 2017) and shown to dynamically decorate the late Golgi, also coalescing after Brefeldin A treatment (Schultzhaus et al., 2017). Given that AP-1 was shown here to associate with late-Golgi, we tested whether it also associates with clathrin light and/or heavy chains, despite having a truncated C-terminal region (Martzoukou et al., 2017).

Figures 4A and 4B suggest a high degree of co-localization of AP-1^σ^ with both clathrin light chain, ClaL, and heavy chain, ClaH. In the case of ClaL, co-migrating foci are often detected with AP-1^σ^, which once formed, move to all dimensions coherently (see also Video 3). In the case of ClaH, “horseshoe”-like structures appear to coalesce predominantly, which again are characterized by coherent movement with AP-1^σ^ (see also Video 4).

**Figure 4:**
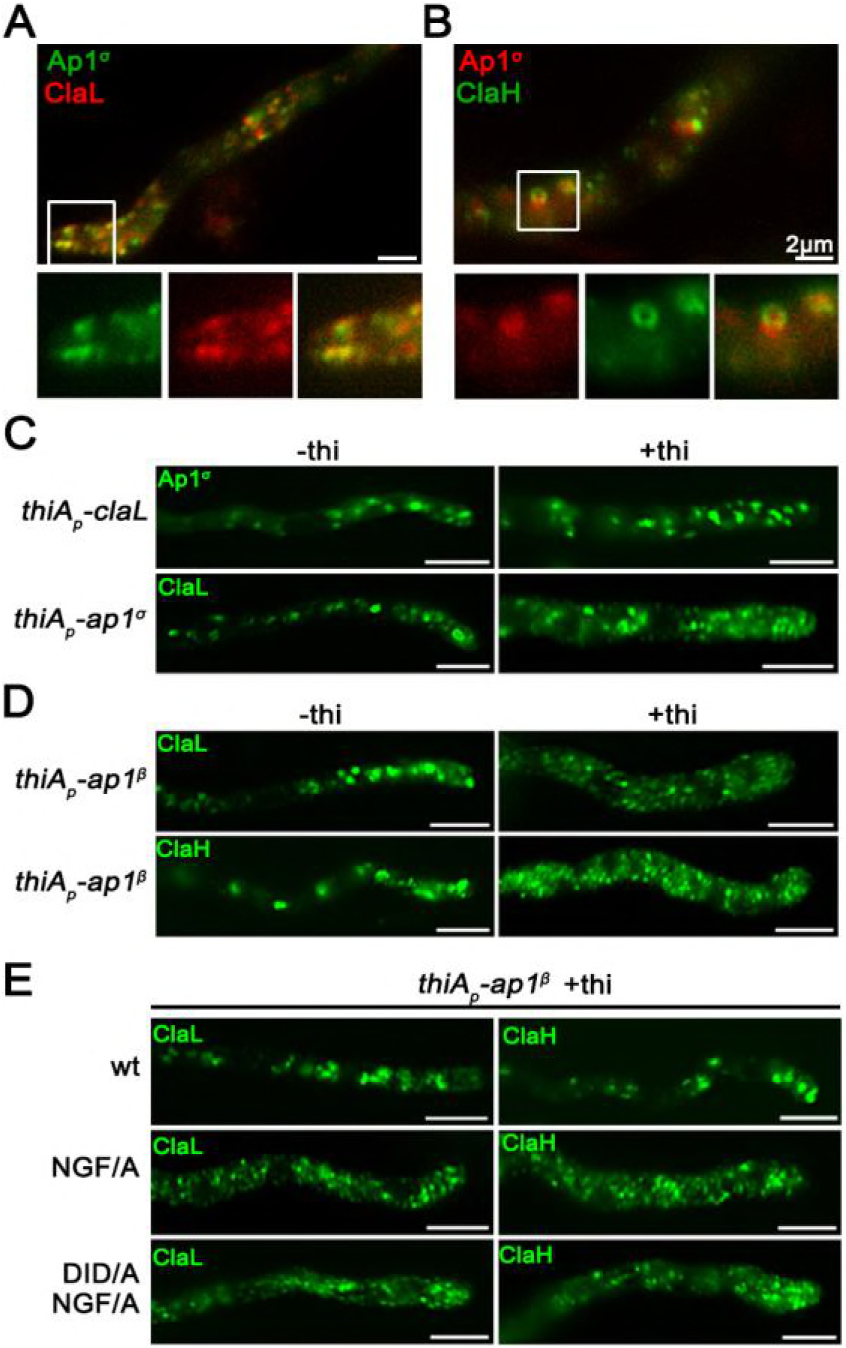
C-terminal motifs in AP-1^β^ are essential for wild-type clathrin localization. (**A, B**) Subcellular localization of Ap1^σ^ relative to that of clathrin light (ClaL) and heavy (ClaH) chains. Notice the significant co-localization of AP-1 with both clathrin chains, also highlighted by the co-migration of the two markers in Videos 3 and 4. (**C, D**) Subcellular distribution of Ap1^σ^ and clathrin light chain ClaL under conditions where *claL* or *ap1^σ^* are repressed, respectively (+thi). Notice that repression of ClaL expression has no significant effect on Ap1^σ^-GFP localization, whereas repression of Ap1^σ^ expression leads to more diffuse ClaL fluorescence with parallel appearance of increased numbers of cytoplasmic puncta. A similar picture is obtained when clathrin light and heavy chain localization are monitored under conditions of expression / repression of *ap1^β^*. These results are compatible with the idea that clathrin localization is dependent on the presence of AP-1, but not vice versa. (**E**) Effect of Ap1^β^ C-terminal mutations modifying putative clathrin binding motifs (^709^NGF/A^711^ and ^632^DID/A^634^) on ClaL and ClaH distribution. Notice that replacement of ^709^NGF^711^, and to a lesser extend of ^632^DID^634^, by alanines, leads to modification of clathrin subcellular localization, practically identical to the picture observed in (**D**) when Ap1^β^ expression is fully repressed. Unless otherwise stated, scale bars represent 5 μm.

We also followed the localization of clathrin in the absence of AP-1, and vice versa, the localization of AP-1 in the absence of clathrin. Results in Figure 4C (upper panel) show that repression of ClaL expression does not affect the wild-type localization of AP-1^σ^. In contrast, repression of AP-1^σ^ leads to a prominent increase in rather static, ClaL-containing, cytoplasmic puncta (Figure 4C, lower panel). This suggests AP-1 functions upstream from ClaL, in line with the established role of AP-1 in clathrin recruitment after cargo binding at late-Golgi or endosomal membranes.

Since our results supported a physical and/or functional association of AP-1 with clathrin, we addressed how this could be achieved given that the *A. nidulans* AP- 1^β^, which is the subunit that binds clathrin in metazoa and yeast (Gallusser and Kirchhausen, 1993), lacks canonical clathrin binding domains in its C-terminal region, but still possesses putative clathrin boxes (^630^LLDID^634^ and ^707^LLNGF^711^). Noticeably, these motifs resemble the LLDLF or LLDFD sequences, found at the extreme C-terminus of yeast AP-1^β^, which have been shown to interact with clathrin (Yeung and Payne, 2001). First, we showed that total repression of AP-1^β^ expression leads to a prominent increase in static ClaL or ClaH puncta, compatible with altered clathrin localization (Figure 4D, upper panels). Then we asked whether the putative clathrin boxes in AP-1^β^ play a role in the proper localization of clathrin. To do so, the ^707^LLNGF^711^ or/and ^630^LLDID^634^ motifs of AP-1^β^ were mutated in a genetic background that also possesses a wild-type *ap-1*^β^ allele expressed via the repressible *thiA*_*p*_ promoter. These strains allowed us to follow the localization of clathrin (*claL* or *claH*) when wild-type or mutant versions of *ap-1*^β^ were expressed. Figure 4D (lower panels) shows that mutations in ^707^LLNGF^711^, and to a much lesser extent ^630^LLDID^634^, lead to modification of clathrin localization, similarly to the picture obtained under total repression of AP-1^β^. Notably, the mutated versions of AP-1^β^ partially restore the growth defects of repressed AP-1^β^ (Figure 4 Supplement 1). This suggests that interaction with clathrin via these boxes is not the primary determinant for the essentiality of AP-1 in fungal growth.

### AP-1 associates with RabE^Rab11^-labeled secretory vesicles

The results obtained thus far suggested that the AP-1 complex is involved in post-Golgi anterograde trafficking of secretory vesicles. In *A. nidulans*, such vesicles deriving from the late-Golgi, traffic along microtubule tracks towards regulated discharge at the apical plasma membrane level (Berepiki et al., 2011; Peñalva et al., 2017; Steinberg et al., 2017; Zhou et al., 2018). Pivotal role in these early processes plays the small GTPase RabE^Rab11^, which recruited along with its regulators, precedes and very probably mediates late-Golgi exit of secretory vesicles towards the hyphal tip (Pantazopoulou et al., 2014; Pinar et al., 2015; Peñalva et al., 2017). Post-Golgi RabE labeled structures, including the Spk, do not co-localize with endosomes stained by FM4-64 or late-endosome/vacuoles stained by CMAC (Figure 5A, 5B). In contrast, they show a significant degree of association with AP-1^σ^, suggesting that the majority of these vesicles are coated by AP-1 (Figure 5C). This is particularly prominent on foci of subapical regions (Figure 5C, right panels) and also at sites of newly emerging branches (Figure 5C, left panels).

**Figure 5:**
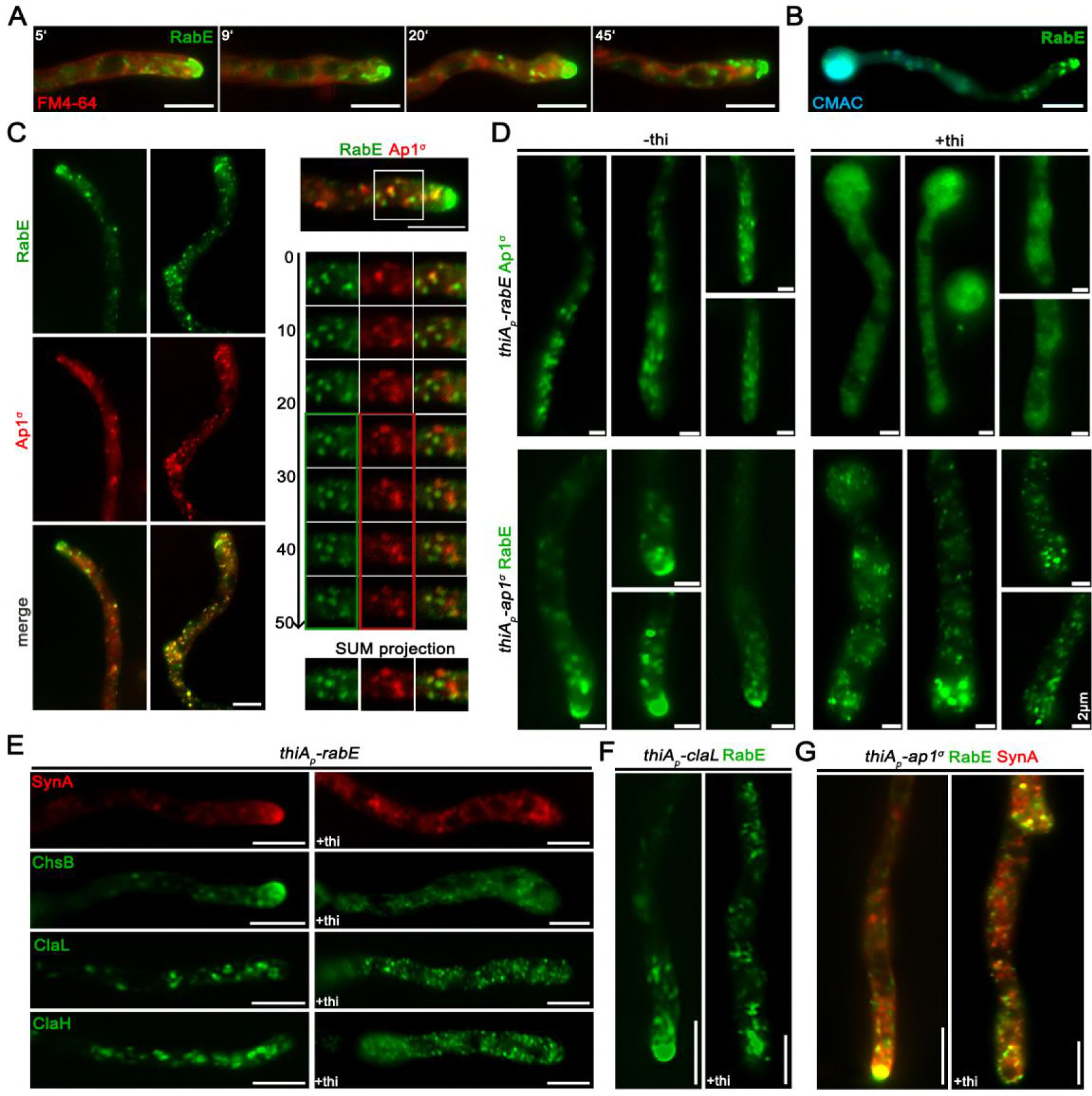
AP-1 associates with RabE^Rab11^-labeled secretory vesicles. (**A, B**) Time course of RabE-GFP localization in the presence of FM4-64 or CMAC, indicating the non-endocytic character for RabE labeled structures. (**C**) Subcellular localization of Ap1^σ^–mRFP and RabE-GFP, showing significant co-localization in several fluorescent cytoplasmic puncta throughout the hyphae but more prominent at sub-apical regions and sites of branch emergence. Notice that co-localization is apparently excluded at the level of Spk, where RabE is prominent, whereas Ap1^σ^ is not. (**D**) Subcellular localization of Ap1^σ^–GFP or RabE-GFP in strains carrying thiamine-repressible *thiA_p_-rabE* or *thiA_p_-ap1^σ^* alleles respectively, observed under conditions of expression (-thi) or repression (+thi). Notice that in the absence of *rabE* expression Ap1^σ^-labeled fluorescence appears as a cytoplasmic haze rather than distinct puncta (upper panels), while in the absence of *ap1^σ^* expression, RabE fluorescence disappears from the Spk and is associated with numerous scattered bright puncta along the hypha (lower panels - 43.24% uniform distribution and intensity of puncta, 37.84% more than two brighter puncta close to the apex are observed, 18.9% one brighter mislocalized punctum at the apex is observed, n=37). (**E**) Subcellular localization of SynA, ChsB, ClaL and ClaH in strains carrying the thiamine-repressible *thiA_p_-rabE* allele, observed under conditions of expression (-thi) or repression (+thi) of *rabE*. Notice that in all cases the wild-type distribution of fluorescence is severely affected, resulting in loss of polarized structures and appearance of an increased number of scattered bright foci, the latter being more evident in ClaL and ClaH. (**F**) Localization of RabE-GFP in a strain carrying a thiamine-repressible *thiA_p_-claL* allele, observed under conditions of expression (-thi) or repression (+thi) of *claL*. Notice the disappearance of RabE from the Spk and its association with numerous scattered bright clusters along the hypha, a picture similar to that obtained in absence of *ap1^σ^* expression in (C). (**G**) Co-localization analysis of SynA and RabE in a strain carrying a thiamine-repressible *thiA_p_-ap1^σ^* allele. Notice that when Ap1^σ^ is expressed, SynA and RabE co-localize intensively at the Spk but also elsewhere along the hypha, whereas when Ap1^σ^ expression is repressed (+thi), both fluorescent signals disappear from the Spk and appear mostly in numerous scattered and rather immotile puncta, several of which show double fluorescence. Unless otherwise stated, scale bars represent 5 μm.

We further examined the association of AP-1 and RabE by following their localization in relevant knockdown mutants. Given that the knockout of RabE proved lethal (results not shown), we monitored AP-1 localization in a knockdown strain where *rabE* expression can be totally repressed via the *thiA*_*p*_ promoter. Similarly, we followed RabE localization in an analogous AP-1 knockdown mutant. Figure 5D (upper panel) shows that when RabE is fully repressed AP-1 fluorescence appears mostly as a cytoplasmic haze, suggesting that AP-1 acts downstream of RabE. Contrastingly, when AP-1 is repressed, RabE does not reach the Spk, while most fluorescence dissolves into scattered static puncta (Figure 5D, lower panel). This strongly suggests that polar secretion of RabE and apparently of secretory vesicles is blocked.

Furthermore, upon repression of *rabE*, crucial apical markers like SynA or ChsB lose their polar distribution, failing to reach their proper destination at the cell cortex (Figure 5E, upper panels). This inhibition of targeting appears more dramatic than the one observed when AP-1 is repressed (see Figure 2), thus indicating the existence of possible alternative RabE-dependent, but AP-1 independent routes. In addition, clathrin labeled structures also lose their wild-type distribution under *rabE* repression conditions, resulting in scattered small puncta (Figure 5E, lower panels), resembling the phenotype observed for ClaL in the absence of a functional AP-1 complex (see Figure 4C, 4D). Similar polar localization defects are observed in RabE- labeled secretory vesicles in strains repressed for clathrin light chain, suggesting that the majority of secretory vesicles requires a clathrin coat to reach the Spk (Figure 5F). Given the fact that, unlike RabE, neither AP-1^σ^ nor clathrin appear to occupy the Spk, it seems that secretory vesicles are uncoated from AP-1 and clathrin prior to their localization in Spk, and thus before actin-dependent localization at the apical PM. We also tested the relative localization of an apical marker (SynA) and RabE in a genetic background where Ap1^σ^ expression can be repressed. When Ap1^σ^ is expressed, SynA and RabE co-localize significantly, mostly evident in the Spk, whereas when Ap1^σ^ is repressed, co-localization persists but shows a more dispersed pattern and is practically absent from of Spk (Figure 5G). This strongly suggests the AP-1 is essential for anterograde movement of post-Golgi vesicles.

### AP-1 associates with the microtubule cytoskeleton

Previous studies have shown that RabE-labeled secretory vesicles utilize microtubule tracks and kinesin-1 for their anterograde traffic, and when present at the Spk use myosin-5 and actin cables to be delivered at the apical PM or eventually move back in retrograde direction powered by dynein motors (Zhang et al., 2011; Egan et al., 2012; Peñalva et al., 2017; Steinberg et al., 2017; Zhou et al., 2018). Here, we examined the possible association of AP-1 with specific dynamic elements of the cytoskeleton involved in cargo traffic. Figure 6A shows that AP-1 puncta decorate microtubules labeled by alpha-tubulin, TubA. Noticeably, the path of motile AP-1 puncta is in most cases dictated by the direction of the microtubules. The association with the microtubule network is further supported by the effect of the anti-microtubule drug Benomyl, which results in an almost complete, but reversible, disassembly of microtubules with a parallel increase in Ap1^σ^-GFP labeled cytoplasmic haze (Figure 6B, upper panel, mostly evident at 4-6 min). In contrast, inhibition of F-actin dynamics via Latrunculin B treatment shows that actin depolymerization does not lead to detectable modification of AP-1 localization (Figure 6B lower panel). This result is in agreement with the observation that AP-1 is excluded from the actin polymerization area.

**Figure 6:**
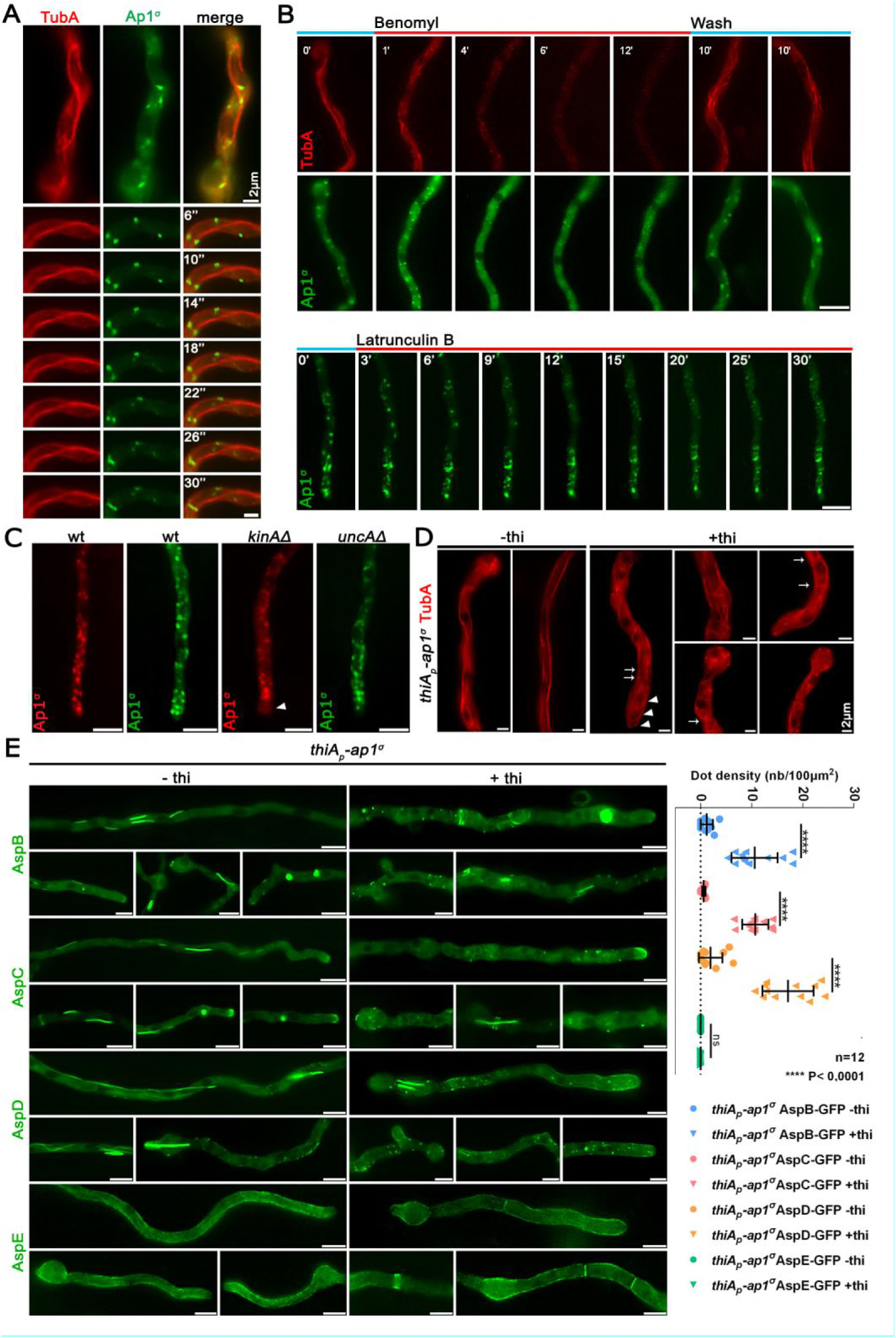
AP-1 associates with the cytoskeleton and affects septin organization. (**A**) Relative Ap1^σ^-GFP and mCherry-TubA (α-tubulin) subcellular localization. Notice Ap1^σ^ fluorescent foci decorating dynamically TubA-labeled microtubules, as highlighted in the selected time lapse images on the lower panels. (**B**) Time course of treatment of strains expressing Ap1^σ^ and TubA with the anti-microtubule drug Benomyl (upper panels). Notice that Benomyl elicits an almost complete, but reversible, disassociation of Ap1^σ^ and TubA, resulting in diffuse cytoplasmic florescent signals. Contrastingly, treatment with the anti-actin drug Latrunculin B does not elicit a significant change in the polar distribution of Ap1^σ^ (lower panels). (**C**) Subcellular localization of AP-1 in wt and in strains lacking the kinesins KinA and UncA, respectively. Notice the absence of apical labeling of AP-1 in the *kinAΔ* strain, indicated with an arrowhead. (**D**) Subcellular organization of the microtubule network, as revealed by TubA-labeling, in a strain carrying a thiamine-repressible *thiA_p_-ap1^σ^* allele, observed under conditions of expression (-thi) or repression (+thi) of *ap1^σ^*. Notice that the absence of Ap1^σ^ leads to a less orientated network, bearing vertical and curved microtubules, and in some cases the appearance of bright cortical spots (2-7 puncta/hypha, usually exhibiting perinuclear localization). (**E**) Subcellular localization of GFP-tagged versions of septins AspB, AspC, AspD and AspE in a strain carrying a thiamine-repressible *thiA_p_-ap1^σ^* allele, observed under conditions of expression (-thi) or repression (+thi) of *ap1^σ^*. Notice that when *ap1^σ^* is repressed, AspB, AspC and AspD form less higher order structures (HOS) such as filaments or bars (*ap1^+^*: 1.58 HOS/hypha, n=87, *ap1^−^*: 0.96 HOS/hypha, n=103) and instead label more cortical spots (see left panel for quantification), some of which appear as opposite pairs at both sides of the plasma membrane, resembling septum formation initiation areas. In contrast, AspE localization remains apparently unaffected under *ap1^σ^* repression conditions. Unless otherwise stated, scale bars represent 5 μm.

Kinesins are motor proteins involved in the transport of secretory vesicles, early endosomes, organelles and also mRNA and dynein motors (Egan et al., 2012; Steinberg 2011; Bauman et al., 2012; Salogiannis and Reck-Peterson, 2017). Based on previous results showing that kinesin-1 KinA (Konzack et al., 2005; Zekert and Fischer, 2009) is the main motor responsible for anterograde traffic of RabE-labeled secretory vesicles, whereas kinesin-3 UncA has no significant role SV secretion (Peñalva et al., 2017), we tested whether KinA and UncA are involved in powering the motility of AP-1 on microtubules. The use of strains carrying deletions of KinA and UncA showed that the motility of AP-1 on microtubules is principally powered by KinA, the absence of which leads to a re-distribution and apparent “stalling” of Ap1^σ^- labeled foci at subapical regions, excluding localization at the hyphal tip area (Figure 6C). This picture is practically identical with the localization of apical cargoes, such as ChsB, in the absence of KinA (Takeshita et al., 2015). In the case of UncA, the Ap1^σ^-labeled foci appear to be largely unaffected, however more prominent localization at the level of Spk and also rather lateral accumulation of relative foci is observed (55,2% of n=25 hyphae) (Figure 6C). These results suggest that UncA might have auxiliary roles in the anterograde traffic of Ap1-labeled secretory vesicles.

The functional association of AP-1 with the cytoskeleton was also investigated by following the appearance of microtubules in a strain lacking AP-1. Figure 6D shows that repression of AP-1 expression led to prominent changes in the microtubule network, as monitored by TubA-GFP fluorescence. These include more curved microtubules towards the apex, distinct bright spots at the periphery of the hyphal head and increased cross sections throughout the hypha, all together suggesting a possible continuous polymerization at the plus end and a problematic interaction with actin through cell-end markers (Takeshita et al., 2013; 2014; Zhou et al., 2018).

### AP-1 is critical for septin organization

Given the role of AP-1 in microtubule organization, we also studied its role on septin localization. Septins are less well characterized GTP-binding proteins, which form hetero-polymers associating into higher order structures, and are thought to play a central role in the spatial regulation and coordination of the actin and microtubule networks in most eukaryotes (Mostowy and Cossart, 2012; Spiliotis, 2018). In *A. nidulans*, five septins have been under investigation, the four core septins AspA-D, which form hetero-polymers appearing in various shapes, including spots, rings and filaments, and a fifth septin of currently unknown function, AspE, not involved in the hetero-polymer and appearing as dense cortical spots at the proximity of the plasma membrane (Hernadez Rodriguez and Momany, 2012; Hernadez Rodriguez et al., 2014; Momany and Talbot, 2017). Figure 6E shows that upon AP-1 repression, hetero-polymer forming core septins AspB, AspC and AspD appear less in the form of filamentous structures, while distinct bright cortical spots tend to accumulate at the hyphal periphery, several of which possibly mark positions of new septa, in agreement with increased numbers of septa observed in the absence of AP-1. Interestingly, AspE, appears largely unaffected with the exception of the more frequent appearance of septa. All the above observations are in agreement with many other previously described phenotypes associating with AP-1 repression and suggest an implication of AP-1 in the processes regulating septin polymer formation. Noticeably, proper endosomal trafficking of septins at growth poles is necessary for growth in *Ustilago maydis* (Bauman et al., 2014).

### AP-1 is involved in endosome recycling

The AP-1 complex has also been implicated in anterograde and retrograde traffic between endosomal compartments and the plasma membrane (Robinson 2004; 2015). However, the existence of relative sorting or recycling endosome, originating from early endosomes (EE), has not been shown rigorously in filamentous fungi (Steinberg et al., 2017).

Major determinants of EE identity are the Rab5 GTPases (Nielsen et al., 1999). The *A. nidulans* Rab5 paralogues RabA and RabB both localize to early endosomes moving on microtubule tracks, with RabB appearing also in relatively static late endosomes. Importantly, the RabA and RabE markers do not co-localize with RabE, which confirms that motile, anterograde-moving, secretory vesicles and motile endosomes are distinct entities (Pantazopoulou et al., 2014). Here, we investigated whether AP-1 associates with Rab5 endosomes.

Figure 7A shows that AP-1 exhibits a degree of transient co-migration with RabB. The coalescence of fluorescence is mostly observed in ring-like structures, which tend to accumulate and convert to more compact forms, suggesting an involvement of AP-1 in recycling, without excluding an additional involvement in vacuolar degradation. Importantly, knockdown of AP-1 led to increased numbers of both RabA and RabB-labeled endosomes (Figure 7B), the majority of which are immotile. In fact, the motile subpopulation of endosomes appears unaffected (Figure 7C). In the absence of AP-1, several distinct RabB foci were also stained by CMAC (Figure 7D), indicating that they are mini-vacuoles, resembling the phenotype of RabA/B in the absence of RabS^Rab7^, a mediator of vacuolar degradation (Abenza et al., 2012). In summary, all evidence presented above strongly support that AP-1 is involved primarily in endosome recycling to the PM, and consequently in its absence, recycling endosomes seem to increase and eventually acquire an identity of degradative endosomes (see also Figure 7E).

**Figure 7:**
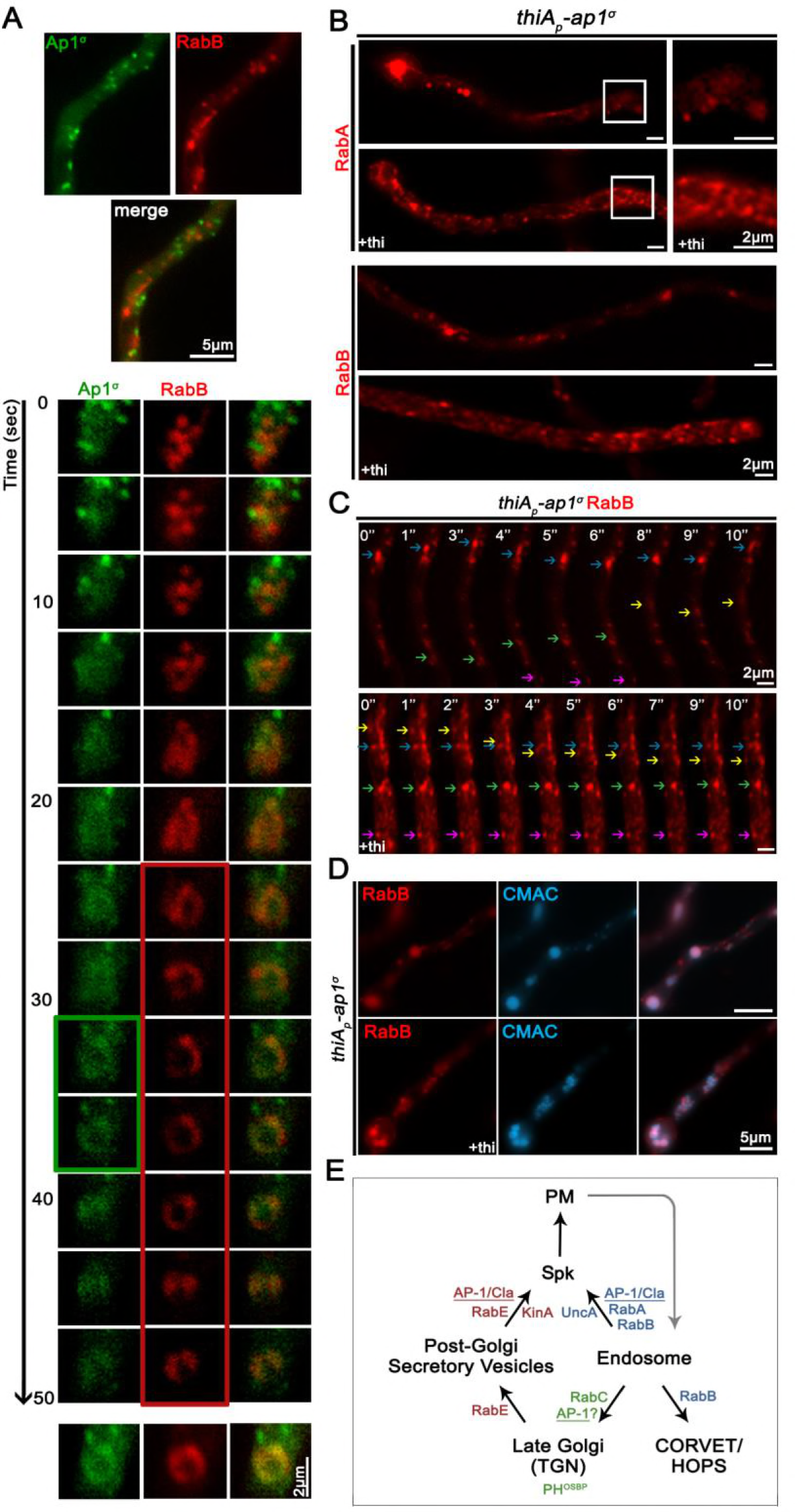
AP-1 is involved in endosome recycling. (**A**) Selected time-lapse images showing the relative localization of Ap1^σ^ and RabB (early endosomal marker). Notice the dynamic association of AP-1 with RabB. (**B**) Subcellular localization of RabA (upper panels) and RabB (lower panels) in a strain carrying a thiamine-repressible *thiA_p_-ap1^σ^* allele, observed under conditions of expression (-thi) or repression (+thi) of *ap1^σ^.* Notice the increased numbers and clustering of both endosomal markers in rather immotile puncta when AP-1 expression is repressed. (**C**) Selected time-lapse images of RabB in a strain carrying a thiamine-repressible *thiA_p_-ap1^σ^* allele, showing that the immotile RabB foci increase in number when *ap1^σ^* is repressed (+thi). However, faster trafficking endosomes can still be observed, in both retrograde and anterograde direction. (**D**) Expression of RabB in a strain carrying a thiamine-repressible *thiA_p_-ap1^σ^* allele, stained with CMAC. Notice that when *ap1^σ^* expression is repressed (+thi), most immotile RabB puncta are stained with CMAC.(**E**) Working model summarizing major findings on the role of the AP-1 complex. Unless otherwise stated, scale bars represent 5 μm.

## Discussion

We have previously shown that the AP-2 complex of *A. nidulans* and probably other higher fungi have a clathrin-independent role in the endocytosis of cargoes necessary for apical recycling of plasma membrane and cell wall components, and thus for fungal polar growth maintenance. This was rather unexpected due to the generally accepted view that AP-2 functions uniquely as a cargo-clathrin adaptor, but also due to its compromised role in the growth of unicellular fungi. Thus, it seems that sorting and trafficking mechanisms are genetically and/or physiologically adaptable in order to meet the specific growth or homeostatic strategies different cells face. In the present work we functionally analyzed the AP-1 complex of *A. nidulans*, as a prototypic example of a simple eukaryote that exhibits continuous polar growth, and showed that AP-1 is indeed essential for cell survival and growth, in a way similar to metazoan cells (Bonifacino, 2014) and probably plants (Robinson and Pimpl, 2013). To our knowledge, no previous study has addressed the role of the AP-1 in filamentous fungi.

In yeasts, which do not maintain polar growth and where the microtubule cytoskeleton is not critical for cargo traffic, AP-1 null mutants are viable, showing relatively moderate growth defects, which in some cases are associated with problematic traffic of specific cargoes, such as chitin synthase Chs3 (Valdivia et al. 2002; Ma et al., 2009; Yu et al., 2013; Arcones et al., 2016). Yeast AP-1 null mutants also have minor defects in lipid PtdIns(3,5)P2-dependent processes and show reduced ability to traffic ubiquitylated cargoes to the vacuole lumen (Phelan et al., 2006). Notably, in *S. cerevisiae*, there are two forms of AP-1 which share the same large (Apl2 and Apl4) and small (Aps1) subunits, but distinct medium subunits (Apm1 or Apm2) that seem to confer differential cargo recognition and sorting (Valdivia et al., 2002; Renard et al., 2010; Whitfield et al., 2016). Additionally, in yeast, the AP-1 complex seems to co-operate with the exomer, a non-essential, fungal-specific heterotetrameric complex assembled at the trans-Golgi network, for the delivery of a distinct set of proteins to the plasma membrane (Hoya et al., 2017; Anton et al., 2018).

Contrastingly to yeasts, repression of AP-1 expression in *A. nidulans* leads to lack of growth, which is related to its inability to maintain apical sorting of all polar cargoes tested, including those necessary for plasma membrane and cell wall biosynthesis. Thus, not only the growth phenotype, but also several underlying cellular defects in AP-1 null mutants resemble those obtained previously with AP-2 loss-of-function mutants (Martzoukou et al., 2017). This is in perfect agreement with the notion that growth of filamentous fungi, unlike yeasts, requires polar apical exocytosis combined with subapical endocytosis and recycling to the apex of specific cargoes related to plasma membrane and cell wall modification (Taheri-Talesh et al., 2008; Peñalva 2010; Shaw et al., 2011). Overall, results presented herein emphasize important differences in membrane trafficking mechanisms employed by yeasts and filamentous fungi, the latter proving a unique genetic and cellular system to dissect cargo sorting in cells characterized by membrane polarity.

Interestingly, despite the similarity in AP-2 and AP-1 phenotypic growth defects, AP-2 has been shown to act independently of clathrin at the PM, while AP-1 is shown here to associate and function with clathrin at several post-Golgi membrane trafficking steps. The similarity of effects caused by null mutations in AP-1 and clathrin chains, concerning RabE^Rab11^-labeled secretory vesicle anterograde traffic and RabA/B^Rab5^-labeled endosome recycling, constitutes strong evidence that AP-1 function is clathrin-dependent. Interestingly, however, the β subunit of AP-1 of *A. nidulans* and all higher fungi lacks the C-terminal appendage domain that contributes to clathrin-binding (Martzoukou et al., 2017). Here, we identified specific short motifs in the C-terminal region of AP-1^β^ that proved critical for proper clathrin subcellular localization and AP-1 function. These motifs (LLNGF and LLDID) resemble motifs shown previously to bind clathrin in yeast (Yeung and Payne, 2001). Thus, contrastingly to the fact that clathrin is dispensable for the function of AP-2 in polar cargo endocytosis it is essential for AP-1-driven polar exocytosis.

A novel point of this work concerns the interaction of AP-1 with RabE^Rab11^. To our knowledge, such an interaction has only been described in a single report in mammalian cells (Parmar et al., 2016). In this case, Rab11 and AP-1 co-localize with the reptilian reovirus p14 FAST protein at the TGN. In metazoa, Rab11 acts as a molecular switch essential for building the necessary molecular machinery for membrane cargo trafficking to the cell surface via its localization and action at the trans-Golgi network, post-Golgi vesicles and specialized recycling endosomes (Welz et al., 2014). In *A. nidulans*, the Rab11 homologue RabE has been previously shown to mark similar subcellular compartments (e.g. late-Golgi and secretory vesicles) and to be involved in anterograde moving of cargoes to the Spk and eventually to the apical PM. Notably, however, RabE does not co-localize with RabA/B^Rab5^-labeled endosomes. The present work strongly suggests that AP-1 and clathrin are sequentially recruited on cargoes, after RabE-dependent maturation of late-Golgi membranes to pre-secretory vesicles, and that secreted cargoes travel embedded within AP-1/clathrin-coated vesicle carriers on MT (see later) to the Spk. At the Spk, AP-1/clathrin coat is most likely released, but RabE remains until the involvement of actin in the last step of fusion with the apical PM.

The impressive similarity of the *A. nidulans* trafficking mechanisms with those of higher organisms is also reflected in the absolute need for proper microtubule (MT) cytoskeleton organization and dynamics (Fischer et al., 2008; Takeshita et al., 2014). We showed that AP-1 is essential for MT organization and associates with microtubules, mainly via KinA. Thus, a specific kinesin motor provides the molecular link between cargo/AP-1/clathrin complexes and cytoskeletal tracks. This is very similar to what has been found in mammalian epithelial cells, where the molecular motor kinesin KIF13A connects AP-1 coated secretory vesicles containing mannose- 6-phosphate receptor to microtubule tracks, and thus mediates their transfer from the TGN to the plasma membrane (Nakagawa et al., 2000). Similarly, in HeLa cells another motor protein kinesin, KIF5, links TGN-derived endosomal vesicles via a direct interaction with Gadkin, a γ-BAR membrane accessory protein of the AP-1 complex, with the microtubule cytoskeleton (Schmidt et al., 2009). Thus, tripartite complexes, including transmembrane cargoes, coat adaptors and motor kinesins, seem to constitute an evolutionary conserved molecular machinery for membrane protein subcellular transport in eukaryotes.

One simple explanation for the essentiality of AP-1 in proper MT organization would be that, in its absence, membrane-associated polarity markers, such as Rho GTPases TeaA or TeaR, which are necessary for microtubule attachment to actin (Fischer et al., 2008; Takeshita and Fischer, 2011; Takeshita et al., 2013; Takeshita 2018), are not sorted correctly in the apex of growing hyphae. Lack of such cell-end markers is known to result in curved or zigzagged organization of MTs and less straight hyphae, compatible with the picture we obtained in the AP-1 null mutant. Importantly, we further supported the essential role of AP-1 in MT organization and function by showing the dramatic effect of the absence of AP-1 on the subcellular organization of septins, proteins that play fundamental roles in the ability of diverse fungi to undergo shape changes and organize the cytoskeleton for polar growth (Zhang et al., 2017; Momany and Talbot, 2017).

Another notable finding of this work concerns the association of AP-1 with recycling endosomes, which represent a pathway distinct from that of RabE-labeled secretory vesicles. Thus, it seems that the combined action of two independent pathways serves the polar distribution of specific cargoes. In *A. nidulans*, early endosomes (EEs) marked by the homologues of the Rab5 family (RabA and RabB) are generated via endocytosis and are easily distinguishable due to their high and long-distance bidirectional motility (Abenza et al., 2009; Steinberg, 2014). A fraction of EEs matures to less motile late endosomes or Multi-Vesicular Bodies (concurrent with increased replacing of RabA/B with RabS^Rab7^), which eventually fuse with vacuoles for cargo degradation (Abenza et al., 2010; 2012; Steinberg, 2014). Another fraction of EEs, mostly the one localized at the subapical collar region of hyphae where very active endocytosis takes place, apparently recycles back to the Spk and from there vesicular cargoes reach the PM (Steinberg, 2014). Whether this takes place directly or *via* retrograde transport to the late-Golgi and anterograde transport in secretory vesicles, is not clear and might well depend on the nature of the cargoes studied. Here we showed that lack of AP-1 leads to a dramatic increase in non-motile RabA/B endosomes, very probably reflecting enhanced maturation into Multi-Vesicular Body endosomes, which suggests that AP-1 has a critical role in the fueling of recycling endosomes to the PM or the late-Golgi. Thus, a consequence of the lack of AP-1 function is compatible with the dramatic increase in static and larger endosomes observed. Similarly, lack of AP-1 function in mammalian cells leads to problematic maturation of early endosomes, associated with aberrant recycling in synapses (Candiello et al., 2016).

Establishing the essential role of AP-1 in polar secretion of specific cargoes in *A. nidulans* (for a schematic view of our findings see Figure 7E and Figure 8), which will probably hold true for other filamentous fungi, also opens a novel little-studied issue. How specific non-polar cargoes are sorted to the plasma membrane? For example, here and previously, we showed that AP-1 and AP-2 complexes are redundant for the proper subcellular expression of transporters that are homogenously present in the PM of growing hyphae and which do not show any indication of polar localization. A critical question to answer is which route(s) and mechanism(s) transporters, and possibly other non-polar transmembrane cargoes (i.e. channels and receptors), use for their sorting, endocytosis or recycling. This question also concerns metazoan and plant cells, where non-polar sorting remains largely understudied. Finally, under the light of previous results obtained in yeast, metazoa or plants, our present work highlights the importance of using different model organisms to address common but evolutionary adaptable mechanisms for membrane cargo traffic in eukaryotes.

**Figure 8:**
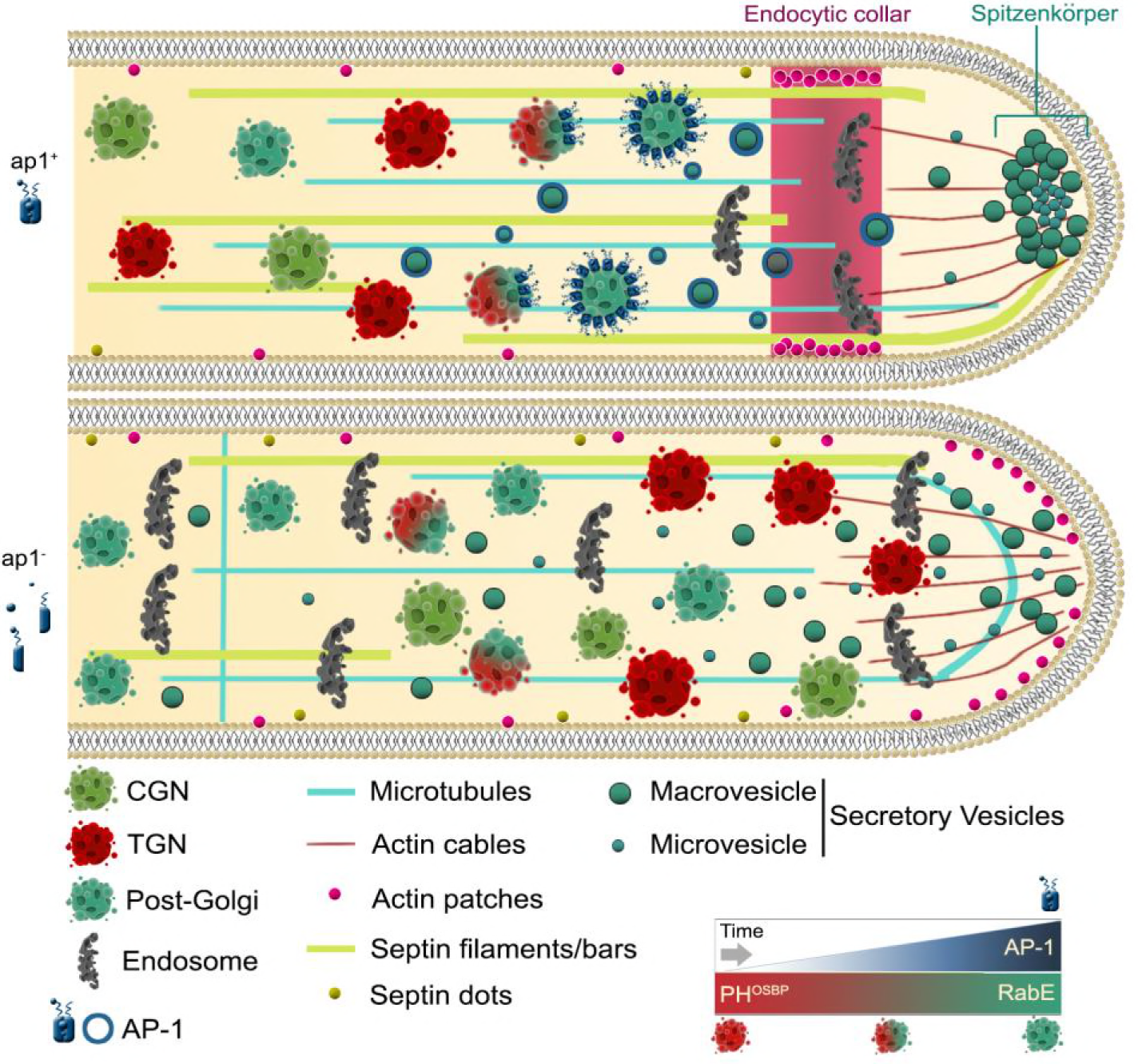
Highly speculative scheme on the role of AP-1 in *A. nidulans* hyphal tip growth.

## Materials and methods

### Media, strains, growth conditions and transformation

Standard complete and minimal media for *A. nidulans* were used (details in FGSC, http://www.fgsc.net.). Media and chemical reagents were obtained from Sigma-Aldrich (Life Science Chemilab SA, Hellas) or AppliChem (Bioline Scientific SA, Hellas). Glucose 0.1-1 % (w/v) was used as a carbon source. NaNO_3_ and NH_4_^+^ (Ammonium tartrate dibasic) were used as nitrogen sources at 10 mM. Thiamine hydrochloride (thi) was used at a final concentration of 10 μM. Transformation was performed as described previously in Koukaki et al. (2003), using an *nkuA* DNA helicase deficient (TNO2A7; Nayak et al., 2006) recipient strain or derivatives for “in locus” integrations of gene fusions, or deletion cassettes by the *A. fumigatus* markers orotidine-5‟-phosphate-decarboxylase (AF*pyrG*, Afu2g0836), GTP-cyclohydrolase II (AF*riboB*, Afu1g13300) and a pyridoxine biosynthesis gene (AFpyroA, Afu5g08090), resulting in complementation of auxotrophies for uracil/uridine (*pyrG89*), riboflavin (*riboB2*) or pyridoxine (*pyroA4*), respectively. Transformants were verified by PCR and Southern analysis. Combinations of mutations and tagged strains with fluorescent epitopes, were generated by standard genetic crossing. *E. coli* strains used were dΗ5α. *A. nidulans* strains used in this study are listed in Supplementary Table 1.

### Nucleic acid manipulations and plasmid constructions

Genomic DNA extraction from *A. nidulans* was performed as described in FGSC (http://www.fgsc.net). Plasmid preparation and DNA gel extraction were performed using the Nucleospin Plasmid kit and the Nucleospin Extract II kit (Macherey-Nagel, Lab Supplies Scientific SA, Hellas). Restriction enzymes were from Takara Bio (Lab Supplies Scientific SA, Hellas). DNA sequences were determined by Eurofins-Genomics (Vienna, Austria). Southern blot analysis using specific gene probes was performed as described in Sambrook et al. (1989), using [^32^P]-dCTP labeled molecules prepared by a random hexanucleotide primer kit and purified on MicroSpin™ S-200 HR columns (Roche Diagnostics, Hellas). Labeled [^32^P]-dCTP (3000 Ci mmol^−1^) was purchased from the Institute of Isotops Co. Ltd, Miklós, Hungary. Conventional PCR reactions, high fidelity amplifications and site-directed mutagenesis were performed using KAPA Taq DNA and Kapa HiFi polymerases (Kapa Biosystems, Roche Diagnostics, Hellas). Gene fusion cassettes were generated by one step ligations or sequential cloning of the relevant fragments in the plasmids pBluescript SKII, or pGEM-T using oligonucleotides carrying additional restriction sites. These plasmids were used as templates to amplify the relevant linear cassettes by PCR. For *ap1^β^* site directed mutations the relevant gene was cloned in the pBS-argB plasmid (Vlanti and Diallinas, 2008). For primers see Supplementary Table 2.

### Protein extraction and western blots

Cultures for total protein extraction were grown in minimal media supplemented with NaNO_3_ or NH_4_^+^ at 25° C. Total protein extraction was performed as previously described (Papadaki et al., 2017). Total proteins (30-50 μg estimated by Bradford assays) were separated in a polyacrylamide gel (8-10 % w/v) onto PVDF membranes (Macherey-Nagel, Lab Supplies Scientific SA, Hellas). Immunodetection was performed with a primary anti-FLAG M2 monoclonal antibody (Sigma-Aldrich), an anti-actin monoclonal (C4) antibody (MP Biomedicals Europe) and a secondary HRP-linked antibody (Cell Signaling Technology Inc, Bioline Scientific SA, Hellas). Blots were developed using the LumiSensor Chemiluminescent HRP Substrate kit (Genscript USA, Lab Supplies Scientific SA, Hellas) and SuperRX Fuji medical X-Ray films (FujiFILM Europe).

### Microscopy and Statistical Analysis

Samples for wide-field epifluorescence microscopy were prepared as previously described (Martzoukou et al., 2017). Germlings were incubated in sterile 35mm μ-dishes, high glass bottom (*ibidi*, Germany) in liquid minimal media for 16-22 h at 25° C. Benomyl, Latrunculin B, Brefeldin A and Calcofluor white were used at final concentrations of 2.5μg ml^−1^, 100μg ml^−1^, 100μg ml^−1^, 0,001% (w/v), respectively. FM4-64 and CMAC staining was according to Peñalva (2005) and Evangelinos et al. (2016), respectively. Images were obtained using a Zeiss Axio Observer Z1/Axio Cam HR R3 camera. Contrast adjustment, area selection and color combining were made using the Zen lite 2012 software. Sum Intensity Projections of selected frames were created using the “Z project” command of ImageJ software. ImageJ Plot profile was used for measurements of fluorescence intensity (https://imagej.nih.gov/ij/). For quantifying dot density in Figure 6, ROIs were selected using the Area Selection tool and the Spot Detector plugin of ICY (http://icy.bioimageanalysis.org/). Tukey‟s Multiple Comparison test was performed (One-way ANOVA) using the Graphpad Prism software for the statistical analysis. Confidence interval was set to 95%. Scale bars were added using the FigureJ plugin of the ImageJ software. Images were further processed and annotated in Adobe Photoshop CS4 Extended version 11.0.2.

## Acknowledgments

We thank Reinhard Fischer for the tagged tubulin, chitin synthase and kinesins strains, Michele Momany for the tagged septins strains and Spyros Efthimiopoulos for the Anti-FLAG M2 antibody. The current research was supported by the *Fondation Santé*, to which we are grateful. OM is suported through the Action “Doctorate Scholarships Programs by the State Scholarships Foundation” by IKY and the European Social Fund (ESF), in the framework of the Operational Program “Education and Life Long Learning” within the NSRF 2014-2020 of the ESF.

## Author contributions

OM, Data curation, Software, Formal analysis, Investigation, Methodology, Conceptualization; GD, Conceptualization, Resources, Formal analysis, Funding acquisition, Validation, Visualization, Manuscript Writing, Project administration; SA, Data curation, Investigation, Methodology, Conceptualization; Manuscript Writing.

## Author ORCIDs

George Diallinas, http://orcid.org/0000-0002-3426-726X

---

**Figure 2 Figure Supplement 1:**
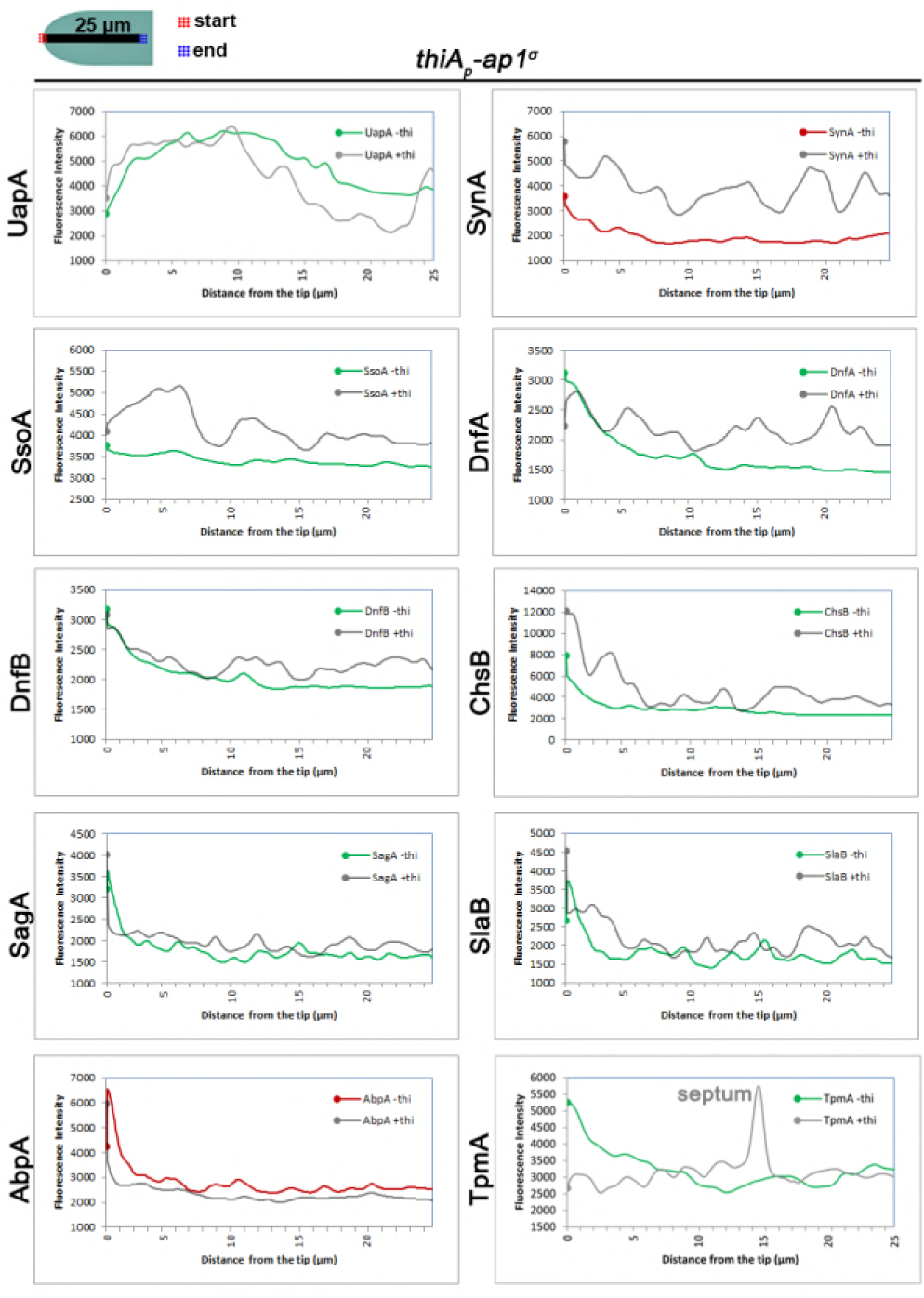
Quantitative analysis of fluorescence intensity of strains shown in Figure 2A, under *ap1^σ^* expressed or fully repressed conditions (-thi, +thi respectively) along 25 μm of hyphal tips. The region measured is depicted in the cartoon on the top left. For details of fluorescence intensity measurements see Materials and methods.

**Figure 2 Figure Supplement 2:**
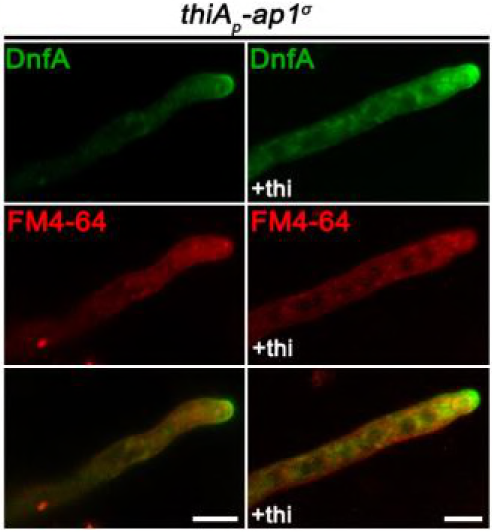
Co-localization of DnfA-GFP with the endocytic dye FM4-64 (10min) indicating that most immotile internal structures are not co-stained with FM4-64.

**Figure 3 Figure Supplement 3:**
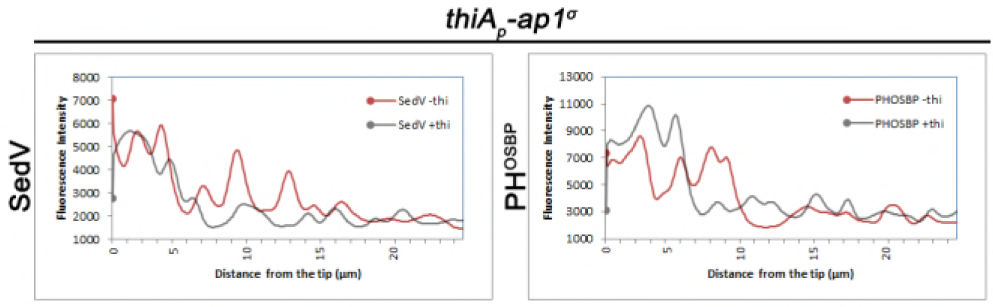
Quantitative analysis of fluorescence intensity of strains shown in Figure 3E, under *ap1^σ^* expressed or fully repressed conditions (-thi, +thi respectively) along 25 μm of hyphal tips. For details of fluorescence intensity measurements see Materials and methods.

**Figure 4 Figure Supplemen 4:**
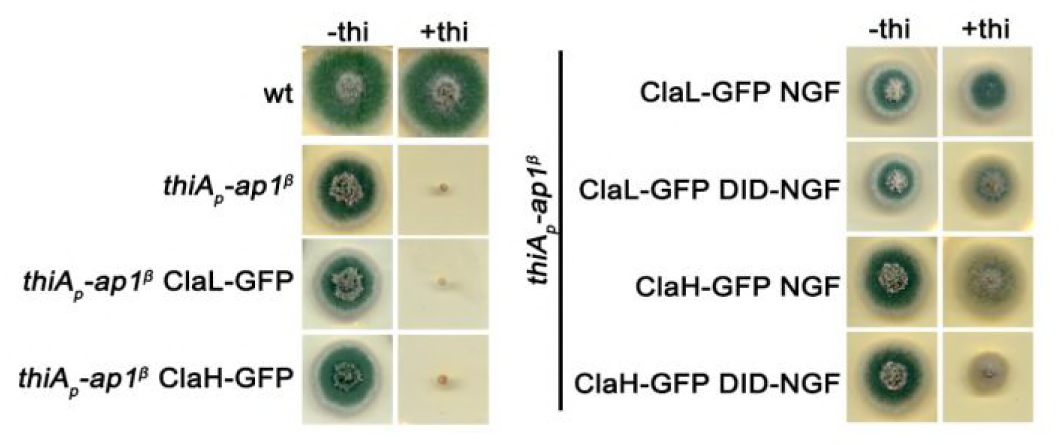
Left panel: Growth test of a standard wild-type (*wt*), a strain carrying a thiamine-repressible *thi*_*p*_-*ap1^β^* allele, and strains expressing ClaL- GFP and ClaH-GFP in the repressible *thi*_*p*_-*ap1^β^* background. Right panel: Growth test of strains carrying the repressible *thi*_*p*_-*ap1^β^* allele “in locus”, together with wt or mutated versions of Ap1^β^ expressed from plasmid integration events, as well as, ClaL-GFP and ClaH-GFP alleles. Notice that expression of the mutated Ap1^β^ versions, which seem defective for clathrin recruitment, partially rescue growth when *thi*_*p*_-*ap1^β^* allele is repressed. This, together with results presented in Figure 4, indicates that total lack of growth observed in the absence of AP-1 is not simply due defective interaction of AP-1 with clathrin.

**Supplementary Table 1:** Strains used in this study. All strains carry the *veA1* mutation affecting sporulation. *pabaA1*, *pyroA4*, *riboB2*, *argB2*, *pyrG89, pantoB100, biA1, nicA2* and *inoB2* are auxotrophic mutations for p-aminobenzoic acid, pyridoxine, riboflavin, arginine, uracil/uridine, D-pantothenic acid, nicotinic acid, biotin and inositol respectively. *yA2* and *wA4* are mutations resulting in yellow and white conidiospore colors respectively.

| Name | Genotype | Reference |
| --- | --- | --- |
| TNO2A7 | *nkuAΔ::argB pyrG89 pyroA4 riboB2* | 8 |
| H1-mRFP | *H1-mRFP::AFriboB nkuAΔ::argB pyrG89 pyroA4* | 9 |
| mRFP-PH<sup>OSBP</sup> | *pyroA4[pyroA::gpdA<sup>m</sup><sub>p</sub>::mRFP-PH<sup>OSBP</sup>] inoB2 niiA4 wA4* | 10 |
| mCherry-sedV | *pyroA4[pyroA<sup>mut</sup>::gpdA<sup>m</sup><sub>p</sub>::mcherry-sedV] nkuAΔ::bar, wA4, niiA4 inoB2* | 11 |
| sagA-GFP | *sagA<sup>(5xGA)</sup>GFP::AFpyrG nkuAΔ::argB pyroA4 riboB2 pyrG89* | 4 |
| slaB-GFP | *slaB-GFP::AFpyrG nkuAΔ::argB pyrG89 pyroA4 argB2* | 2 |
| mCherry-synA<br>GFP-tpmA | *mCherry-synA::AFpyrG yA::AFpyroA GFP-tpmA fwA1 pyrG89 pyroA4 nicA2 nkuAΔ::argB* | 16 |
| abpA-mRFP | *abpA-mRFP::AFpyrG yA2 pabaA1 pyrG89* | 2 |
| ssoA-GFP | *GFP-ssoA::AFpyrG nkuAΔ::bar pyrG89 pyroA4* | 16 |
| rabOts | *rabI<sup>A136D</sup>::AFpyrG pabaA1* | 12 |
| sedVts | *sedV<sup>R238G</sup>::AFpyrG pyroA4 pyrG89 nkuAΔ::bar* | 12 |
| chsB-GFP | *alcA<sub>p</sub>::GFP-chsB::NCpyr4 nkuAΔ::argB pyrG89 pyroA4* | 17 |
| dnfA-GFP | *dnfA-GFP::AFpyrG nkuAΔ::argB pyrG89 pabaA1 pyroA4* | 13 |
| dnfB-GFP | *dnfB-GFP::AFpyrG nkuAΔ::argB pyrG89 pabaA1 pyroA4* | 13 |
| thiA<sub>p</sub>-ap1<sup>σ</sup> | *thiA<sub>p</sub>::FLAG ap1<sup>σ</sup>::AFriboB nkuAΔ::argB pyrG89 pyroA4 riboB2* | 7 |
| ap2<sup>σ</sup>-mRFP | *ap2<sup>σ</sup>-(5xGA)mRFP::AFpyrG nkuAΔ::argB pyrG89 riboB2 pyroA4* | 7 |
| thiA<sub>p</sub>-claL | *thiA<sub>p</sub>::claL::AFpyrG nkuAΔ::argB pyroA4 riboB2 pyrG89* | 7 |
| thiA<sub>p</sub>-ap1<sup>σ</sup><br>uapA-GFP | *thiA<sub>p</sub>::FLAG ap1<sup>σ</sup>::AFriboB alcA<sub>p</sub>-uapA-GFP pabaA1* | 7 |
| claL-GFP | *claL<sup>(5xGA)</sup>GFP::AFpyrG nkuAΔ::argB pyroA4 riboB2 pyrG89* | 7 |
| claH-GFP | *claH<sup>(5xGA)</sup>GFP::AFpyrG nkuAΔ::argB pyroA4 riboB2 pyrG89* | 7 |
| claL-mRFP | *claL<sup>(5xGA)</sup>mRFP::AFpyrG nkuAΔ::argB pyroA4 riboB2 pyrG89* | 7 |
| thiA<sub>p</sub>-claH<br>dnfA-GFP | *thiA<sub>p</sub>-claH::AFpyroA dnfA-GFP::AFpyrG nkuAΔ::argB pyrG89 pabaA1 pyroA4* | 7 |
| ap2Δ dnfA-GFP | *dnfA-GFP::AFpyrG ap2<sup>σ</sup>Δ::AFriboB nkuAΔ::argB pyrG89 pyroA4* | 7 |
| mCherry-rabA | *alcA<sub>p</sub>::mCherry-rabA::argB yA2 pantoB100 argB2* | 1 |
| mRFP-rabB | *alcA<sub>p</sub>::mRFP-rabB::pyroA niiA4 nkuAΔ::bar inoB2 pyroA4 wA3* | 1 |
| aspB-GFP | *aspB-GFP::AFpyrG pyrG89 argB2 pabaB22 nkuAΔ::argB riboB2* | 19 |
| aspC-GFP | *aspC-GFP::AFpyrG pabaA6 biA1* | 6 |
| aspD-GFP | *aspD-GFP::AFpyrG argB2 riboB2* | 3 |
| aspE-GFP | *aspE-GFP::AFpyrG riboB2* | 3 |
| mCherry-tubA | *alcA<sub>p</sub>::mCherry-tubA::pyroA nkuAΔ::argB pyrG89 pyroA4* | 17 |
| GFP-kinA<sup>rigor</sup> | *kinAΔ::NCpyr4 alcA<sub>p</sub>::GFP-kinA<sup>rigor</sup>::pyroA pyrG89 pyroA4 argB2* | 14, 20 |
| claH-GFP | *claH<sup>-(5xGA)</sup>GFP::AFpyrG nkuAΔ::argB pyrG89 pyroA4 riboB2* | This study |
| ap1<sup>σ</sup>-GFP | *ap1<sup>σ</sup>-(5xGA)<sup>-</sup>GFP::AFpyrG nkuAΔ::argB pyrG89 pyroA4 riboB2* | This study |
| ap1<sup>σ</sup>-mRFP | *ap1<sup>σ</sup>-(5xGA)<sup>-</sup>mRFP::AFpyrG nkuAΔ::argB pyrG89 pyroA4 riboB2* | This study |
| mRFP-rabB<br>ap1<sup>σ</sup>-GFP | *alcA<sub>p</sub>::mRFP-rabB::pyroA4 ap1<sup>σ</sup>-(5xGA)<sup>-</sup>GFP::AFpyrG nkuAΔ::argB* | This study |
| thiA<sub>p</sub>-ap1<sup>σ</sup><br>mCherry-rabA | *thiA<sub>p</sub>::<sup>FLAG</sup>ap1<sup>σ</sup>::AFriboB alcA<sub>p</sub>::mCherry-rabA::argB nkuAΔ::argB pantoB100 pyroA4* | This study |
| thiA<sub>p</sub>-ap1<sup>σ</sup><br>mRFP-rabB | *alcA<sub>p</sub>::mRFP-rabB::pyroA thiA<sub>p</sub>::<sup>FLAG</sup>ap1<sup>σ</sup>::AFriboB nkuAΔ::argB riboB2 pyroA4 wA3* | This study |
| thiA<sub>p</sub>-ap1<sup>σ</sup> GFP-ssoA | *thiA<sub>p</sub>::<sup>FLAG</sup>ap1<sup>σ</sup>::AFriboB GFP-ssoA::AFpyrG pabaA1* | This study |
| thiA<sub>p</sub>-ap1<sup>σ</sup><br>mCherry-sedV | *thiA<sub>p</sub>::<sup>FLAG</sup>ap1<sup>σ</sup>::AFriboB pyroA4::[pyroA<sup>mut</sup>::gpdA<sup>m</sup><sub>p</sub>::mcherry-sedV] nkuAΔ::bar inoB2* | This study |
| ap1<sup>σ</sup>-GFP<br>mCherry-sedV | *ap1<sup>σ</sup>-(5xGA)<sup>-</sup>GFP::AFpyrG pyroA4[pyroA<sup>mut</sup>::gpdA<sup>m</sup><sub>p</sub>::mcherry-sedV] nkuAΔ::argB pyrG89 pyroA4 inoB2* | This study |
| thiA<sub>p</sub>-ap1<sup>σ</sup><br>sagA-GFP | *thiA<sub>p</sub>::<sup>FLAG</sup>ap1<sup>σ</sup>::AFriboB sagA<sup>-(5xGA)</sup>GFP::AFpyrG nkuAΔ::argB pyrG89 pyroA4 riboB2 (4)* | This study |
| thiA<sub>p</sub>-ap1<sup>σ</sup> slaB-GFP | *thiA<sub>p</sub>::<sup>FLAG</sup>ap1<sup>σ</sup>::AFriboB slaB-GFP::AFpyrG nkuAΔ::argB pyrG89 pyroA4 riboB2* | This study |
| thiA<sub>p</sub>-ap1<sup>σ</sup><br>abpA-mRFP | *thiA<sub>p</sub>::<sup>FLAG</sup>ap1<sup>σ</sup>::AFriboB abpA-mRFP::AFpyrG nkuAΔ::argB pyrG89 pyroA4 riboB2* | This study |
| thiA<sub>p</sub>-ap1<sup>σ</sup> H1-mRFP | *thiA<sub>p</sub>::<sup>FLAG</sup>ap1<sup>σ</sup>::AFriboB H1-mRFP::AFriboB* | This study |
| ap1<sup>σ</sup>-GFP<br>thiA<sub>p</sub>-ap1<sup>μ</sup> | *thiA<sub>p</sub>::ap1<sup>μ</sup>::AFriboB ap1<sup>σ</sup>-(5xGA)<sup>-</sup>GFP::AFpyrG nkuAΔ::argB pyrG89 pyroA4 riboB2* | This study |
| ap1<sup>σ</sup>-GFP<br>thiA<sub>p</sub>-ap1<sup>β</sup> | *thiA<sub>p</sub>::ap1<sup>β</sup>::AFriboB ap1<sup>σ</sup>-(5xGA)<sup>-</sup>GFP::AFpyrG nkuAΔ::argB pyrG89 pyroA4 riboB2* | This study |
| thiA<sub>p</sub>-ap1<sup>σ</sup><br>mCherry-tubA | *thiA<sub>p</sub>::<sup>FLAG</sup>ap1<sup>σ</sup>::AFriboB alcA<sub>p</sub>::mCherry-tubA:pyroA* | This study |
| kinA<sup>rigor</sup>-GFP<br>ap1<sup>σ</sup>-mRFP | *kinAΔ:pyr4 pyroA4[alcA<sub>p</sub>::kinA<sup>rigor</sup>-GFP:pyroA] ap1<sup>σ</sup>-(5xGA)<sup>-</sup>mRFP::AFpyrG pyrG89* | This study |
| ap1<sup>σ</sup>-mRFP<br>claH-GFP | *ap1<sup>σ</sup>-(5xGA)<sup>-</sup>mRFP::AFpyrG claH<sup>-(5xGA)</sup>GFP::AFpyrG nkuAΔ::argB pyroA4* | This study |
| ap1<sup>σ</sup>-mRFP<br>dnfA-GFP | *dnfA-GFP::AFpyrG ap1<sup>σ</sup>-(5xGA)<sup>-</sup>mRFP::AFpyrG nkuAΔ::argB pyrG89 pyroA4* | This study |
| ap1<sup>σ</sup>-GFP<br>mCherry-synA | *mCherry-synA::AFpyrG ap1<sup>σ</sup>-(5xGA)<sup>-</sup>GFP::AFpyrG nkuAΔ::argB pyrG89 pyroA4 nicA2* | This study |
| thiA<sub>p</sub>-ap1<sup>σ</sup><br>dnfA-GFP<br>ap2<sup>σ</sup>Δ | *thiA<sub>p</sub>::<sup>FLAG</sup>ap1<sup>σ</sup>::AFriboB ap2<sup>σ</sup>Δ::AFpyroA dnfA-GFP::AFpyrG nkuAΔ::argB pyroA4* | This study |
| thiA<sub>p</sub>-ap1<sup>σ</sup><br>dnfB-GFP | *thiA<sub>p</sub>::<sup>FLAG</sup>ap1<sup>σ</sup>::AFriboB dnfB-GFP::AFpyrG nkuAΔ::argB pyrG89 pyroA4* | This study |
| thiA<sub>p</sub>-ap1<sup>σ</sup><br>claL-GFP | *thiA<sub>p</sub>::<sup>FLAG</sup>ap1<sup>σ</sup>::AFriboB claL<sup>(5xGA)</sup>-GFP::AFpyrG nkuAΔ::argB pyrG89 pyroA4 riboB2* | This study |
| ap1<sup>σ</sup>-GFP<br>mCherry-tubA | *alcA<sub>p</sub>::mCherry-tubA::pyroA ap1<sup>σ</sup>-(5xGA)-GFP::AFpyrG nkuAΔ::argB pyrG89 pyroA4* | This study |
| thiA<sub>p</sub>-rabE | *thiA<sub>p</sub>::rabE::AFriboB nkuAΔ::argB pyrG89 pyroA4 riboB2* | This study |
| thiA<sub>p</sub>-rabE<br>ap1<sup>σ</sup>-GFP | *ap1<sup>σ</sup>-(5xGA)-GFP::AFpyrG thiA<sub>p</sub>::rabE::AFriboB nkuAΔ::argB pyrG89 pyroA4 riboB2* | This study |
| ap1<sup>σ</sup>-GFP<br>thiA<sub>p</sub>-claL | *ap1<sup>σ</sup>-(5xGA)-GFP::AFpyrG thiA<sub>p</sub>::claL::AFriboB nkuAΔ::argB pyrG89 pyroA4 riboB2* | This study |
| thiA<sub>p</sub>-ap1<sup>σ</sup><br>aspB-GFP | *aspB-GFP::AFpyrG thiA<sub>p</sub>::<sup>FLAG</sup>ap1<sup>σ</sup>::AFriboB nkuAΔ::argB pyrG89 riboB2 pyroA4* | This study |
| thiA<sub>p</sub>-ap1<sup>σ</sup><br>aspC-GFP | *aspC-GFP::AFpyrG thiA<sub>p</sub>::<sup>FLAG</sup>ap1<sup>σ</sup>::AFriboB pabaA6 pyroA4* | This study |
| thiA<sub>p</sub>-ap1<sup>σ</sup><br>aspD-GFP | *aspD-GFP::AFpyrG thiA<sub>p</sub>::<sup>FLAG</sup>ap1<sup>σ</sup>::AFriboB riboB2* | This study |
| thiA<sub>p</sub>-ap1<sup>σ</sup><br>aspE-GFP | *aspE-GFP::AFpyrG thiA<sub>p</sub>::<sup>FLAG</sup>ap1<sup>σ</sup>::AFriboB pyroA4* | This study |
| thiA<sub>p</sub>-rabC | *thiA<sub>p</sub>::rabC::AFriboB nkuAΔ::argB pyrG89 pyroA4 riboB2* | This study |
| thiA<sub>p</sub>-rabC<br>ap1<sup>σ</sup>-GFP | *thiA<sub>p</sub>::rabC::AFriboB ap1<sup>σ</sup>-(5xGA)-GFP::AFpyrG nkuAΔ::argB pyrG89 pyroA4 riboB2* | This study |
| GFP-rabE | *GFP-rabE::AFpyrG pyrG89 pyroA4 riboB2 nkuAΔ::argB* | This study |
| GFP-rabE ap1<sup>σ</sup>-<br>mRFP | *ap1<sup>σ</sup>-(5xGA)-mRFP::AFpyrG GFP-rabE::AFpyrG nkuAΔ::argB* | This study |
| thiA<sub>p</sub>-ap1<sup>σ</sup><br>mCherry-synA<br>GFP-tpmA | *thiA<sub>p</sub>::<sup>FLAG</sup>ap1<sup>σ</sup>::AFriboB yA::AFpyroA GFP-tpmA AFpyrG::mCherry-synA nkuAΔ::argB pyrG89* | This study |
| ap1<sup>σ</sup>-GFP<br>mRFP-PH<sup>OSBP</sup> | *ap1<sup>σ</sup>-(5xGA)-GFP::AFpyrG [pyroA-gpdA<sup>m</sup><sub>p</sub>::mRFP-PH<sup>OSBP</sup>]pyroA4 nkuAΔ::argB pyrG89 inoB2* | This study |
| thiA<sub>p</sub>-ap1<sup>σ</sup><br>mRFP-PH<sup>OSBP</sup> | *pyroA4[pyroA-gpdA<sup>m</sup><sub>p</sub>::mRFP-PH<sup>OSBP</sup>] thiA<sub>p</sub>::<sup>FLAG</sup>ap1<sup>σ</sup>::AFriboB* | This study |
| thiA<sub>p</sub>-ap1<sup>σ</sup>ap2<sup>σ</sup>Δ<br>dnfA-GFP | *thiA<sub>p</sub>::<sup>FLAG</sup>ap1<sup>σ</sup>::AFriboB ap2<sup>σ</sup>Δ::AFpyroA dnfA-GFP::AFpyrG nkuAΔ::argB pyroA4* | This study |
| rabO<sup>ts</sup> ap1<sup>σ</sup>-GFP | *rab1<sup>A136D</sup>::AFpyrG ap1<sup>σ</sup>-(5xGA)-GFP::AFpyrG pyroA4* | This study |
| sedV<sup>ts</sup> ap1<sup>σ</sup>-GFP | *sedV<sup>R238G</sup>::AFpyrG ap1<sup>σ</sup>-(5xGA)-GFP::AFpyrG nkuAΔ::bar pyroA4* | This study |
| thiA<sub>p</sub>-ap1<sup>σ</sup> GFP-rabE mCherry-synA | *thiA<sub>p</sub>::<sup>FLAG</sup>ap1<sup>σ</sup>::AFriboB GFP-rabE::AFpyrG AFpyrG::mCherry-synA nkuAΔ::argB pyrG89 pyroA4* | This study |
| ap1<sup>σ</sup>-GFP<br>uncAΔ | *ap1<sup>σ</sup>-(5xGA)-GFP::AFpyrG uncAΔ::AFriboB nkuAΔ::argB pyrG89 riboB2 pyroA4* | This study |
| thiA<sub>p</sub>-rabE claL-GFP | *thiA<sub>p</sub>-rabE::AFriboB claL<sup>-(5xGA)</sup>GFP::AFpyrG nkuAΔ::argB pyrG89 riboB2 pyroA4* | This study |
| thiA<sub>p</sub>-rabE claH-GFP | *thiA<sub>p</sub>-rabE::AFriboB claH<sup>-(5xGA)</sup>GFP::AFpyrG nkuAΔ::argB pyrG89 riboB2 pyroA4* | This study |
| thiA<sub>p</sub>-claL GFP-rabE | *thiA<sub>p</sub>-claL::AFriboB GFP-rabE::AFpyrG nkuAΔ::argB pyrG89 pyroA4 riboB2* | This study |
| thiA<sub>p</sub>-rabE mCherry-synA | *thiA<sub>p</sub>-rabE::AFriboB AFpyrG::mcherry-synA nkuAΔ::argB pyrG89 pyroA4* | This study |
| thiA<sub>p</sub>-rabE GFP-chsB | *thiA<sub>p</sub>-rabE::AFriboB AFpyrG::alcA<sub>p</sub>-GFP-chsB nkuAΔ::argB pyrG89 pyroA4 riboB2* | This study |
| thiA<sub>p</sub>-ap1<sup>β</sup> claH-GFP | *thiA<sub>p</sub>-ap1<sup>β</sup>::AFpyrG claH<sup>-(5xGA)</sup>GFP::AFpyrG argB2* | This study |
| thiA<sub>p</sub>-ap1<sup>β</sup> claL-GFP | *thiA<sub>p</sub>-ap1<sup>β</sup>::AFpyrG claL<sup>-(5xGA)</sup>GFP::AFpyrG argB2* | This study |
| thiA<sub>p</sub>-ap1<sup>β</sup> claH-GFP ap1<sup>β</sup> | *thiA<sub>p</sub>-ap1<sup>β</sup>::AFpyrG claH<sup>-(5xGA)</sup>GFP::AFpyrG ap1<sup>β</sup>::argB argB2* | This study |
| thiA<sub>p</sub>-ap1<sup>β</sup> claL-GFP ap1<sup>β</sup> | *thiA<sub>p</sub>-ap1<sup>β</sup>::AFpyrG claL<sup>-(5xGA)</sup>GFP::AFpyrG ap1<sup>β</sup>::argB argB2* | This study |
| thiA<sub>p</sub>-ap1<sup>β</sup> claH-GFP ap1<sup>β</sup> NGF/A | *thiA<sub>p</sub>-ap1<sup>β</sup>::AFpyrG claH<sup>-(5xGA)</sup>GFP::AFpyrG ap1<sup>β</sup>-N709A/G710A/F711A::argB argB2* | This study |
| thiA<sub>p</sub>-ap1<sup>β</sup> claH-GFP ap1<sup>β</sup> DID/A NGF/A | *thiA<sub>p</sub>-ap1<sup>β</sup>::AFpyrG claH<sup>-(5xGA)</sup>GFP::AFpyrG ap1<sup>β</sup>-D632A/I633A/D634AN709A/G710A/ F711A argB2* | This study |
| thiA<sub>p</sub>-ap1<sup>β</sup> claL-GFP ap1<sup>β</sup> NGF/A | *thiA<sub>p</sub>-ap1<sup>β</sup>::AFpyrG claL<sup>-(5xGA)</sup>GFP::AFpyrG ap1<sup>β</sup>-N709A/G710A/F711A::argB argB2* | This study |
| thiA<sub>p</sub>-ap1<sup>β</sup> claL-GFP ap1<sup>β</sup> DID/A NGF/A | *thiA<sub>p</sub>-ap1<sup>β</sup>::AFpyrG claL<sup>-(5xGA)</sup>GFP::AFpyrG ap1<sup>β</sup>-D632A/I633A/D634AN709A/G710A/ F711A::argB argB2* | This study |

**Supplementary Table 2:**
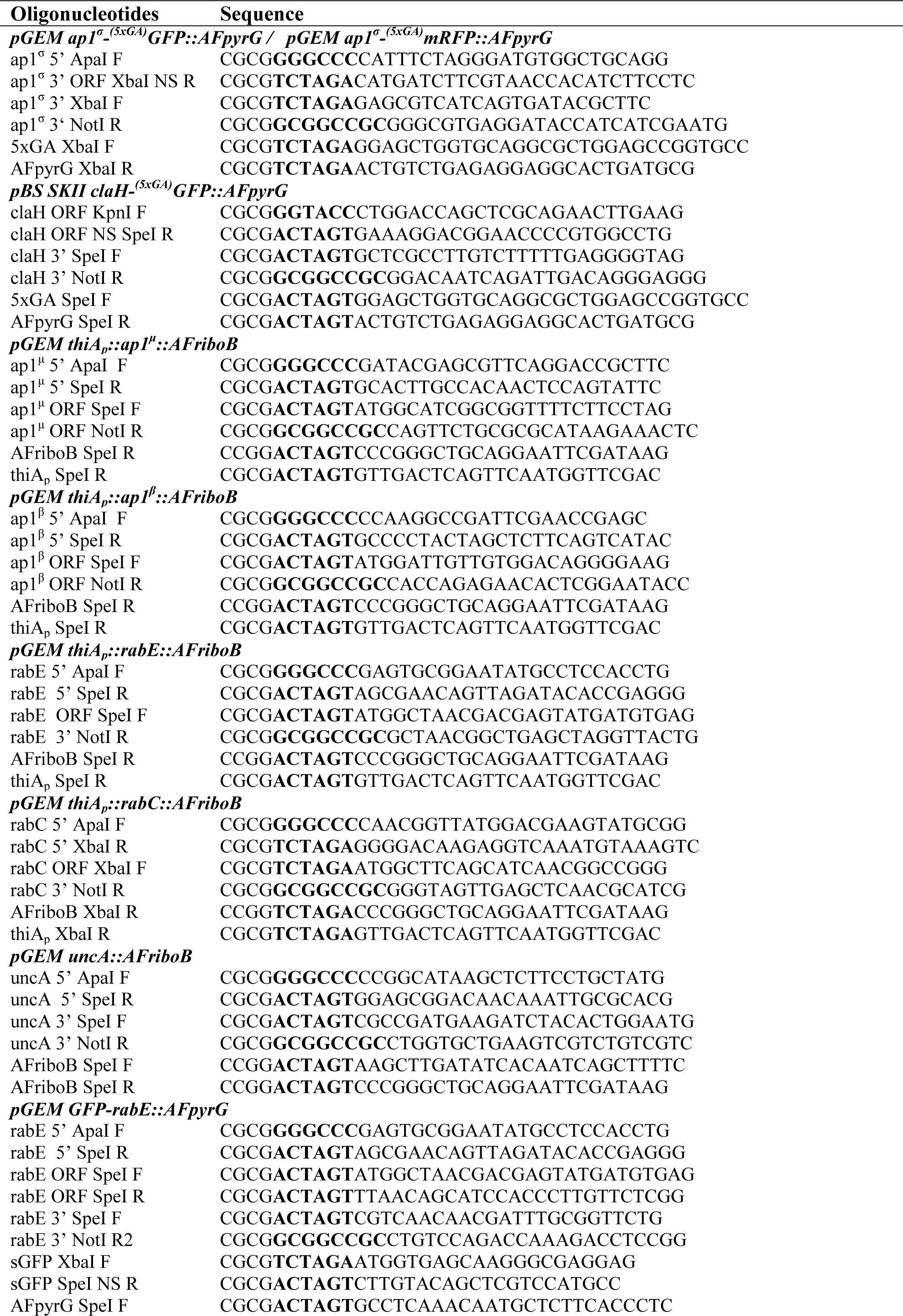
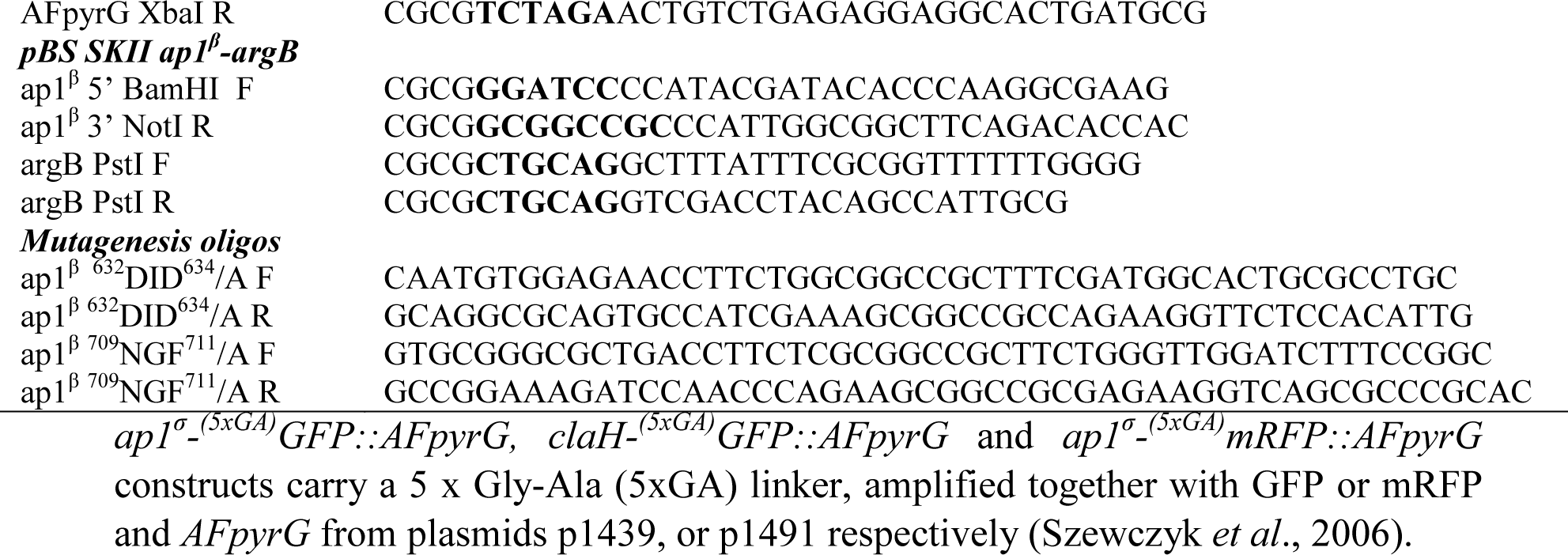
Oligonucleotides used in this study for cloning purposes *ap1*^*σ*^-^*(5xGA)*^*GFP::AFpyrG, claH*-^*(5xGA)*^*GFP::AFpyrG* and *ap1*^*σ*^-^*(5xGA)*^*mRFP::AFpyrG* constructs carry a 5 x Gly-Ala (5xGA) linker, amplified together with GFP or mRFP and *AFpyrG* from plasmids p1439, or p1491 respectively (Szewczyk *et al*., 2006).

